# Hyperdivergent haplotypes control melanic camouflage in a polymorphic moth

**DOI:** 10.64898/2026.02.24.707733

**Authors:** Luca Livraghi, Jordan L. Cassily, Mallory Brayer, Alexander T. Carter, Jasmine D. Alqassar, Sheina B. Sim, Scott M. Geib, Joseph J. Hanly, Arnaud Martin

## Abstract

A majority of moths use melanic pigmentation to blend into the visual environment, and some species display genetically controlled color polymorphisms that provide tractable systems for dissecting the genetic basis of camouflage. Here we leveraged the natural polymorphism of *Anticarsia gemmatalis* moths to map a major-effect gene that controls melanic variation, and study its developmental roles using expression and loss-of-function assays. A genome-wide association mapping identifies the *ivory:mir-193* non-coding RNA locus as a major locus of melanic variation, with highly-divergent haplotypes differentiating the Light and Dark morphs. Second, the expression of *ivory:mir-193* prefigures the melanic coloration of each morph, and suggests that *cis*-regulatory variation at this locus determines the difference in adult pattern. Finally, CRISPR disruption of the *ivory* promoter and *mir-193* hairpin region each abolish melanic scale pigmentation, demonstrating that this lnc-pri-miRNA module is required for dark-scale differentiation. These findings corroborate the emerging role of *ivory:mir-193* as a key factor in the development of melanic patterning, and highlight its unique propensity to repeatedly drive phenotypic variation of adaptive potential in butterflies and moths.

**Highlights:**

- *Anticarsia gemmatalis* display striking variation in camouflage patterns
- Dark vs. Light color polymorphism map to the *ivory:mir-193* hotspot locus
- Highly-diverged haplotypes drive morph-specific expression of *ivory:mir-193*
- The *miR-193* miRNA is required for melanic scale development

**Graphical Abstract:** 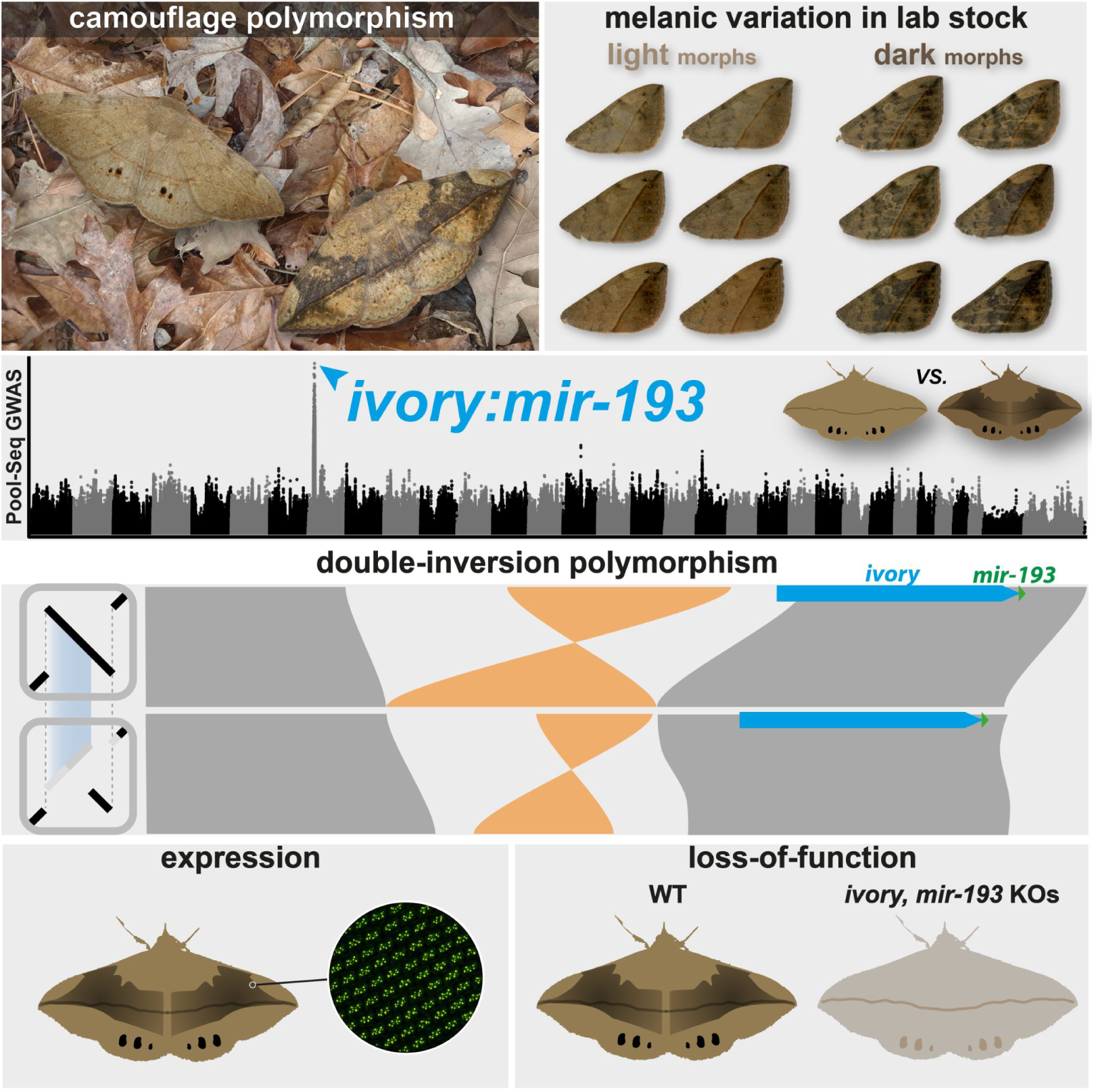

## Introduction

Camouflage represents one of nature’s most striking examples of an organism’s ability to adapt to a specific environment. To hide, animals can match the color and texture of the background where they rest^1^. To achieve this they employ varying strategies, including concealment tricks such as countershading^2^, depth illusion^3^, and edge effects^4^, or directly mimicking elements of the environment that are recognized as inert objects, such as leaves or twigs (often referred to as masquerading^5^). Coloration traits sometimes vary within a given species, and such polymorphisms enable the discovery of genetic loci that drive phenotypic adaptation in natural populations^6,7^. In vertebrates, a few major effect genes underlie the matching of cryptic traits to distinct visual environments^8,9^, such as in populations of deer mice^10,11^ and lizards^12^ that are locally adapted to sand color, or in rabbit and hare populations that grow a white coat in regions with extended snow covering^13,14^. Among insects, the industrial melanism of peppered moths (*Biston betularia*) is an iconic case of rapid adaptation to anthropogenic change^15,16^, where the spread of black morphs tracked the darkening of tree bark due to industrial pollution, and is controlled by a single, large effect locus^17^. Likewise, the ability of Orange Oakleaf butterflies (*Kallima inachus*) to masquerade the color and textural details of dead leaves is orchestrated by several alleles segregating at a single master locus^18^.

Importantly, genetic analyses of camouflage adaptation in butterflies and moths (Lepidoptera) have started to reveal a unifying principle that had been unsuspected before the genomic era; both the peppered moth and the oakleaf butterfly loci map to the same region in the vicinity of the gene *cortex*^17,18^. In fact, that same locus underlies other variations involved in crypsis, aposematism, mimicry, and seasonal polyphenism across lineages of butterflies and moths that span more than 150 MY of divergence^19^. In butterflies, this includes the genera *Heliconius*^20–24^, *Junonia*^25^, *Kallima*^18^, *Hypolimnas*^26^, *Melinaea*, *Mechanitis*, *Hypothiris*^27^, *Papilio*^28^, and *Pieris*^29^. In moth lineages, this includes the genera *Biston*^30,31^, *Phigalia*, *Odontopera*^32^, *Utetheisa*^33^, *Chetone*^27^, and *Ectropis*^34^. This level of repeatability makes this locus a genomic hotspot of color pattern variation, and suggests a predictable genetic basis underpinning the evolution of melanic adaptations in Lepidoptera^7,35,36^.

Recent studies revealed that the non-coding transcript *ivory:mir-193*, rather than *cortex*, is the causal gene regulating these polymorphisms across Lepidoptera. This locus transcribes the *ivory* long non-coding RNA, named after white mutants of *Heliconius melpomene* that carry a 78-kb deletion in that region^37^. The 3′ end of *ivory* lies immediately upstream of the miRNA genes *mir-193* and *mir-2788*, and expression profiling suggests that *mir-193* is processed from the *ivory* primary transcript itself^23,38^. In this configuration, *ivory* functions as a host gene that forms a lnc-pri-miRNA, *ie.* a long non-coding primary transcript involved in the expression of miRNA loci^39,40^. CRISPR evidence in multiple species indicates that the conserved *ivory* promoter region and *mir-193* are both required for melanic patterning, while *cortex* and *mir-2788* are not^24,34,38,41^. Importantly, RNA *in situ* hybridizations of *ivory* transcripts in pupal wings prefigure the exact position of melanic patterns^24,38,41,42^, implying that *ivory* expression integrates regulatory signals that sketch minute details of the adult phenotype. In summary, these data show that *ivory:mir-193* forms a single gene unit, and that it integrates complex spatial information to orchestrate the production of *miR-193* miRNA to modulate scale cell precursor identity and pigmentation.

Here we studied the genetic basis of melanic variation in the velvetbean caterpillar (*Anticarsia gemmatalis*), and show that the *ivory:mir-193* locus is a master regulator of melanic patterning in this polymorphic moth. Pooled genomic sequencing followed by genome-wide association showed that individuals segregating for dark/light pigmentation phenotypes map to a large 600-kb region centered around the *ivory:mir-193* locus. Functional perturbation experiments using CRISPR-Cas9 KOs revealed that both the *ivory* promoter and *mir-193* hairpin are required for melanic scale development, and expression assays further revealed that *ivory* transcription is spatially associated with dark scales in this species. Together, these results position *ivory:mir-193* as a central regulator of melanic pigmentation that can tune both local scale patterning and whole-body melanism, providing a genetic lever for the diversification of camouflage strategies in *A. gemmatalis*.

## Results

### Melanic variation in Light and Dark morphs maps to the *ivory:mir-193* locus

*A. gemmatalis* is a pan-American species of erebid moths (**Fig. 1A**), and a notorious pest of legume crops^43,44^. This species also shows remarkable phenotypic variation in its camouflage patterns, with specimens often differing in hue, darkness, and textural details including stripes, speckling, and mottling (**Fig. 1B**). To gain insight into the genetics of this polymorphism, we reared specimens from a commercially available outbred population (Stoneville strain) and separated moths into two extreme phenotypic categories: “Light” morphs with a homogenous beige, and “Dark” morphs which show melanic mottling across the wing (**Fig. 1C-D, Fig. S1**). Preliminary F_1_ and F_2_ crosses (**Figs. S2–S3**) indicated that the Light versus Dark variation segregates as a Mendelian trait, with Dark representing the dominant form and Light the recessive form (56 Dark : 18 Light F_2_, Chi-square test for 3:1 ratio, P > 0.89). To carry out a genome wide-association study of Light *vs.* Dark variation in this population, we first generated a reference genome from a Dark female individual, resulting in a chromosome-level genome assembly of 391.7 Mb (**Fig. S4**), (NCBI: GCA_050436995.1). We next sequenced one DNA pool each of 23-25 females per Dark/Light phenotypic class (**Fig. 1D**), achieving a normalized median sequencing depth of 88X per pool^45^. We then performed a genome-wide Pool-Seq association scan by applying Fisher’s exact test to compare allele counts between the Dark and Light pools at each SNP. This analysis revealed a single peak of associated variants on chromosome 8, corresponding to 600 kb of sequence immediately upstream of the *ivory:mir-193* locus and spanning 33 annotated genes with no known roles in pigmentation, except for *ivory* exon 1(**Fig. 1E** and **Table S1**). These results implicate the *ivory:mir-193* locus as a major-effect determinant of melanic camouflage variation in *A. gemmatalis*.

**Figure 1.**
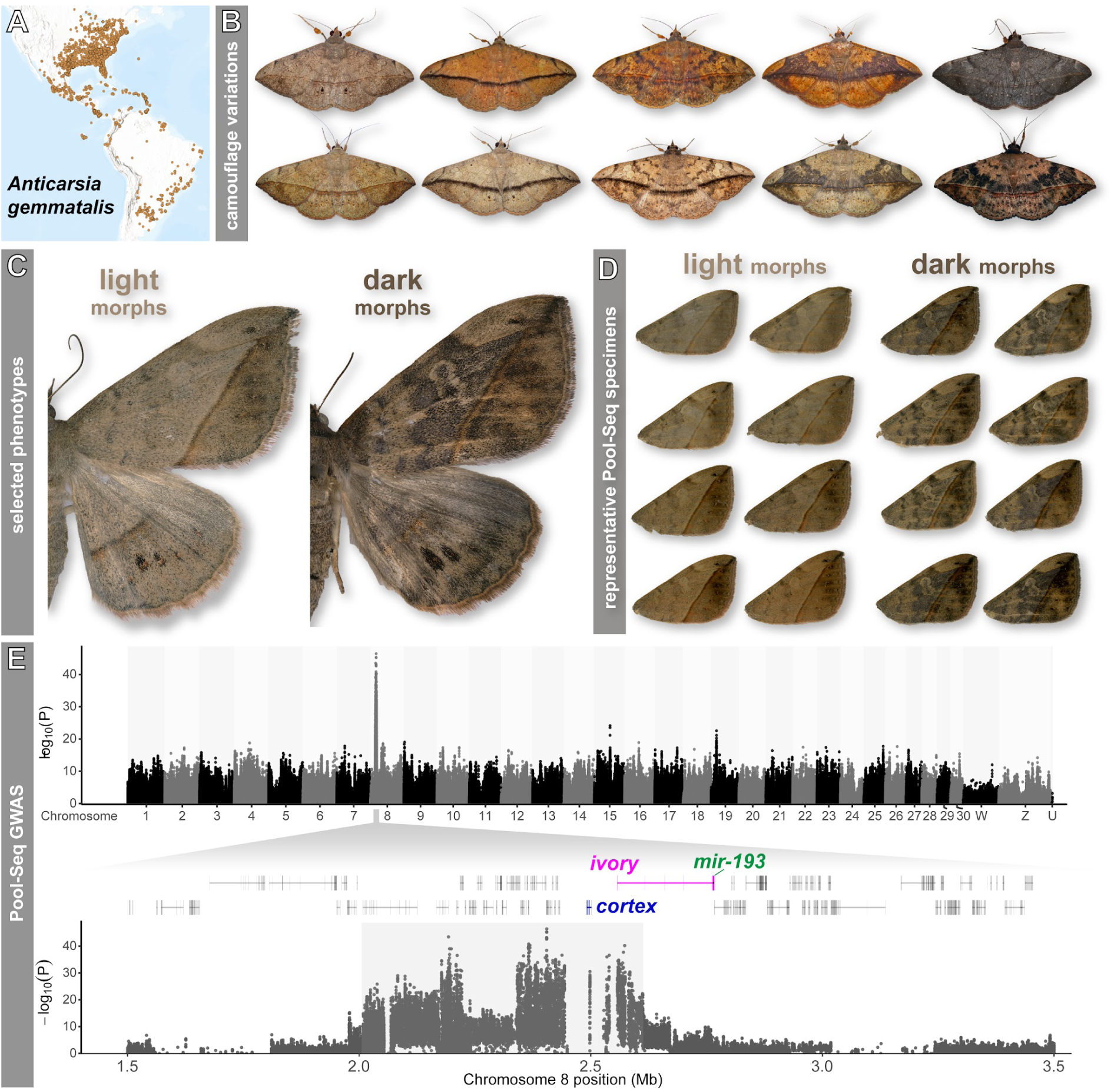
Levels of melanism map to a single major effect locus in the polymorphic moth *A. gemmatalis*. **(A)** *A. gemmatalis* distribution throughout the Americas based on GBIF occurrence data. **(B)** Overview of color morphs in the Eastern US (images retrieved from iNaturalist entries, all CC-BY). **(C)** Representative female Light and Dark morphs segregating through our lab reared population. **(D)** Representative dorsal forewings of the individuals used in the Pool-Seq experiment for each phenotypic class. **(E)** Genome-wide association scan for melanism based on pooled sequencing of Dark and Light morphs. For each SNP, allele counts in the two phenotypic pools were compared using Fisher’s exact test implemented in PoPoolation2, and *–log₁₀(P)* values are plotted by genomic position, revealing a single peak of association on Chromosome 8. Below is a zoomed view of the associated region on Chromosome 8, highlighting SNPs with the strongest association. Gene models in the interval are shown, with the positions of *cortex*, *ivory*, and *mir-193* indicated.

To confirm the presence of *ivory:mir-193* transcripts in developing wing tissues, and to help annotate the locus in detail, we generated both standard and small RNA-seq libraries at 40-80% and 40% of pupal development respectively. Standard RNA-seq libraries revealed a large transcript containing 6 exons, spanning ∼200 kb and terminating close to the site of *mir-193* transcription start (**Fig. S5A**). Small RNA-seq libraries confirmed the presence of both 5’ and 3’ arms of *mir-193* and *mir-2788*, with coverage analysis indicating a 3’ mature and 5’ star strand for *mir-193* (**Fig. S5B**). In summary, differences in melanic pigmentation strongly associate with the *ivory:mir-193* locus, and both transcripts are expressed during pupal wing development in *A. gemmatalis*.

### A divergent haplotype block 5’ of *ivory:mir193* associates with melanic variation

Mapping the Pool-Seq reads to the Dark reference genome revealed a broad reduction in short-read coverage across the associated interval (**Fig. 2A**). This reduction was most pronounced among the Light morphs but also noticeable among Dark morphs, consistent with the presence of Light-recessive and Dark-dominant alleles in each pool. Read mapping dropout is most evident in intergenic and intronic regions (**Fig. S6**). Re-mapping of the Pool-Seq data against a genome assembly from a Light individual reversed this coverage pattern, but changing the reference did not impact the Pool-GWA signal (**Fig. S7**). Consistent with this, F_ST_ and D_XY_ estimates of genetic differentiation indicated substantial sequence divergence between morph-associated haplotypes across the entire GWA interval (**Fig. 2B** and **Fig. S8-9**). When combining all samples, we found higher levels of nucleotide diversity (π) in the GWA interval compared to the rest of the genome (**Fig. 2B** and **Fig. S8-9**), regardless of phenotype. These estimates of F_ST_, D_XY_ and π are concordant with the drop in read-mapping in this region of interest, and point at the existence of large haplotypes underlying melanic variation, rather than a narrow peak driven by few linked causal mutations.

**Figure 2.**
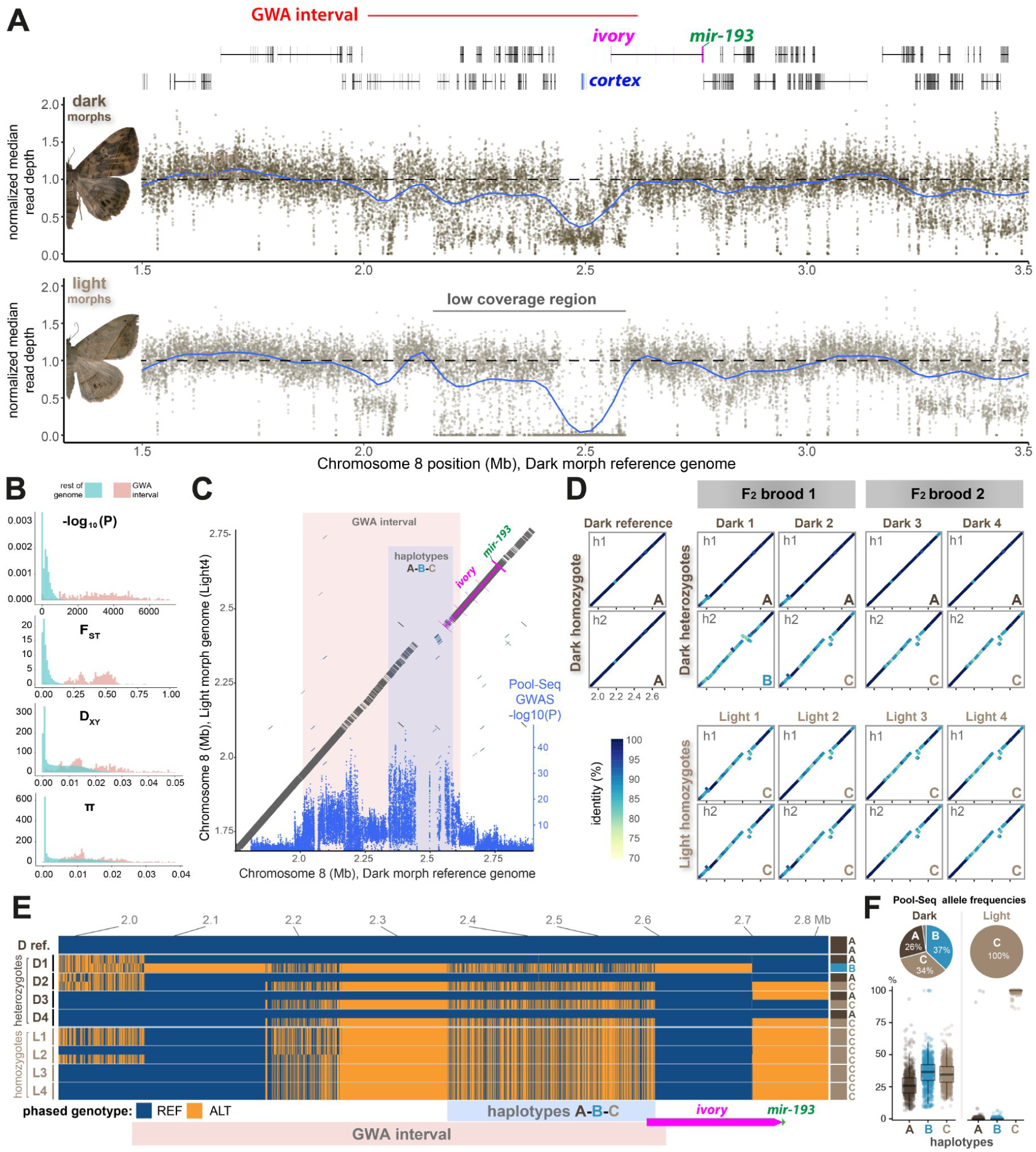
A haplotype block of three divergent haplotypes differentiates Light and Dark morphs. **(A)** Read-depth profiles for the Dark and Light pooled sequencing libraries across the GWA interval, using the Dark reference genome. Dotted line: 1.0x coverage baseline. **(B)** Density plots of F_ST_ and D_XY_ statistics between Dark and Light Pool-Seq groups, and π within the combined Pool-Seq datasets, highlighting the deep divergence of the melanic locus (red) compared to the rest of the genome (blue). **(C)** Combined plot showing GWA SNPs overlaid with a dotplot alignment of reference Dark and Light haplotypes. The GWA interval includes a 430-kb region of hyperdivergence between Light and Dark morphs (as highlighted in panel A), including a large collinear block with spotty alignment, and a region of low similarity new *ivory:mir-193*. **(D)** Sequence dotplots comparing all HiFi phased haplotypes (h1, h2) of 8 F_2_ individuals from two separate broods to the Dark reference assembly, across the associated locus. Colors indicate sequence similarity (%) over 500-bp blocks. **(E)** SNP genotype plot showing variant sites from HiFi phased haplotypes (blue = reference, orange = alternate) across the GWA interval. Dark individuals were heterozygous for the reference-like haplotype A and alternate haplotype C across the hyperdivergent region. Light individuals were homozygous for haplotype C. **(F)** Estimated haplotype frequencies of the A-B-C haplotype groups among the 25 Dark and 23 Light individuals in the Pool-Seq groups. Whisker plots feature the mean frequency of 2,312 B-informative and 2,305 C-informative biallelic SNPs across the A-B-C haplotype block in each Pool-Seq group (**Fig. S11**).

To further resolve the haplotype structure of this locus, we generated PacBio HiFi long-read data from eight F_2_ individuals derived from two independent crosses (2 Dark + 2 Light in each brood), and assembled phased haplotypes across the candidate region. Pairwise comparisons of Dark and Light haplotypes revealed a large stretch of divergent sequence that spans most of the GWA interval, followed by a zone of Structural Variation (SV) immediately 5’ of *ivory:mir193* (**Fig. 2C-D and Fig. S10**). This region includes a 230-kb haplotype block where each allele can be assigned to one of three haplotypes, dubbed A, B, and C **(Fig. 2D-E**). In our limited samples of phased genomes, the five Dark individuals carried haplotype A in homozygous or heterozygous form (A/A, A/B or A/C), whereas the four Light individuals were homozygous for haplotype C. Haplotype B was sampled from a single Dark individual. This pattern of segregation suggests that its three haplotypes may underlie the differences between Dark and Light morphs, as also hinted by the fact that the haplotype block coincides with the region of highest GWA signal (**Fig. 2C**).

Next we estimated the allelic frequencies of the haplotypes A-B-C in the Pool-Seq groups of 25 Dark and 23 Light individuals, using a total of 4,617 diagnostic sites isolated from the 230-kb haplotype block (**Fig. 2F and Fig. S11)**. Haplotype C showed a 100% frequency in the Light pool: all the Light-recessive morphs were C/C homozygotes. In contrast, we found a mixture of haplotypes A (26% mean frequency), B (37%) and C (34%) in the Dark pool. This asymmetric distribution makes haplotypes A and B incompatible with a Light phenotype, as it is unlikely that either of them would have escaped random sampling from a second pool of 23 samples (*P* < 6 x 10^-4^, see STAR★Methods). It follows that both haplotypes A and B cause a dominant Dark effect, and further work will be required to assess their additive or epistatic effects on variation within the Dark morph category (**Fig. 1D**). In summary, these results show that the melanic interval segregates for three deeply divergent haplotypes that extend over 230-kb in the 5’ region of *ivory:mir-193*, with both haplotypes A and B associated with Dark morphs, and haplotype C producing the recessive phenotype of Light morphs.

### A double-inversion differentiates the three haplotypes 5′ of *ivory:mir-193*

We next focused on haplotypes A-B-C to characterize the structural basis of their divergence in more detail. The GWA interval showed a relative enrichment in transposable elements (TE) compared to the rest of the genome (**Fig. 3A and Fig. S12**). However, no substantial differences in TE class representation were observed among the three haplotypes, suggesting that their divergence is not primarily explained by the invasion or amplification of any specific TE class.

**Figure 3.**
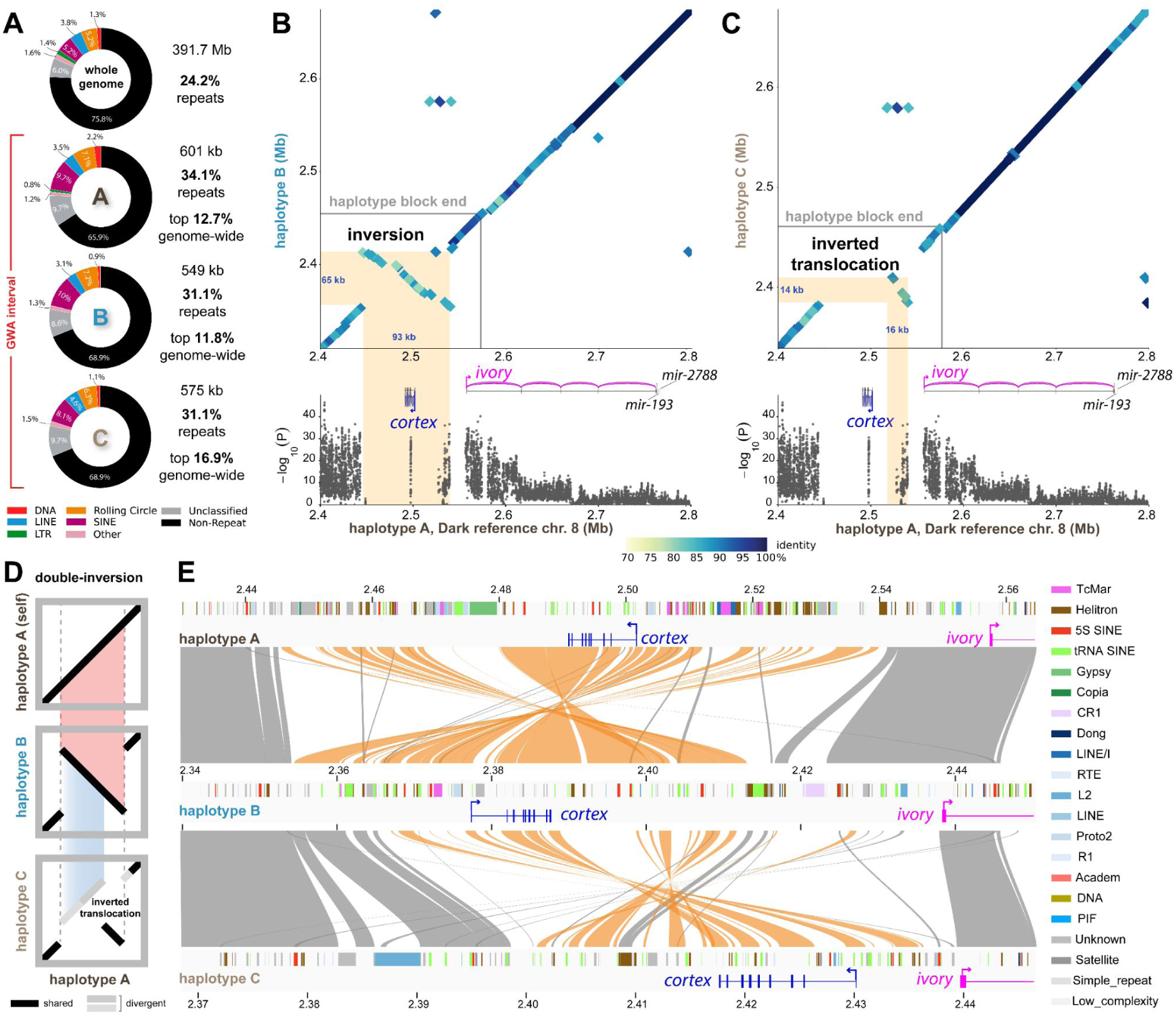
A double-inversion underlies complex structural variation 5’ of *ivory:mir-193.* **(A)** TE class composition across the entire reference genome and the GWA interval, for the 3 haplotypes. The GWA region is enriched in repeat content compared to the rest of the genome (>84.1% percentile among size-matched windows). **(B-C)** Dot plot comparisons of haplotypes B and C relative to the reference genome (haplotype A), Highlights demarks large SVs 5’ of *ivory:mir-193*, including a large inversion (haplotype B) and a translocated inversion (haplotype C). **(D)** Schematic relationship between observed SVs. Haplotypes A and B differ by a large inversion. A nested inversion between B and C underlie the pattern of inverted transposition observed between haplotypes A and C, with additional nucleotide divergence explaining the low Pool-Seq read coverage and drop in GWA signal in this region. **(E)** Ribbon plots of the 3 haplotypes highlighting stepwise differences in microsynteny and TE annotation. Inverted blocks are shown in orange. Collinear blocks and spurious hits in repeated elements are shown in grey.

Pairwise dotplot comparisons revealed that haplotype B differs from the reference-like haplotype A by a large inversion in the 5′ region of *ivory:mir-193* (**Fig. 3B**). Haplotype C, by contrast, showed a more complex relationship to haplotype A, indicating a short inverted transposition nested in a region of high nucleotide divergence (**Fig. 3C**). Importantly, the A-B inversion and A-C inverted translocation share a nearly identical breakpoint (**Fig. S3**), suggesting the inverted translocation is the product of a double inversion^46^. Microsynteny alignments showed that the rearrangements are embedded within a broader block of sequence divergence immediately upstream of *ivory:mir-193* (**Fig. 3D-E and Fig. S13**). These comparisons support a stepwise structural model in which haplotype B arose from A through a large inversion, with haplotype C subsequently arising through a second inversion nested within that rearranged block. Alternatively, haplotype C may be representative of the ancestral state and could have led to B and then A via the reverse path. However, we note that this scenario may require *de novo* regulatory innovation to generate the more complex Dark patterns, and that it is therefore less parsimonious than a loss (in haplotype C) of Dark enhancers pre-existing in the A–B lineage.

A combination of large-scale rearrangement, TE richness, and extensive nucleotide divergence likely explain both the severe Pool-Seq coverage dropout and the uneven distribution of SNP-based association signal across the interval. Thus, morph-associated variation at the *ivory:mir-193* locus is embedded in a structurally complex regulatory landscape comprising at least three large, highly differentiated haplotypes.

### Expression of *ivory:mir-193* prefigures melanic patterning

To further test whether transcriptional regulation of *ivory:mir-193* is associated with melanic scale development in *A. gemmatalis*, we performed RT–qPCR on developing forewing tissue from Dark and Light female morphs across a pupal time course (**Fig. 4A-B**). Expression of *ivory* was strongly stage-dependent: Light individuals showed an earlier transient increase at 30% of pupal development, whereas Dark individuals exhibited a pronounced peak at the 45% stage, at which point *ivory* expression was significantly higher in Dark than Light developing wings (Mann–Whitney U-test, *p* = 0.029). Expression in both morphs converged again at 60% before declining by 75%. By contrast, *miR-193* expression remained low through mid-development but increased sharply at later stages in Dark individuals, with significant morph-specific divergence at 60% of pupal development (Mann–Whitney U-test, *p* = 0.024). This temporal delay relative to the *ivory* lncRNA profile is consistent with miRNA accumulation following transcription of the *ivory* precursor and subsequent processing^38^.

**Figure 4.**
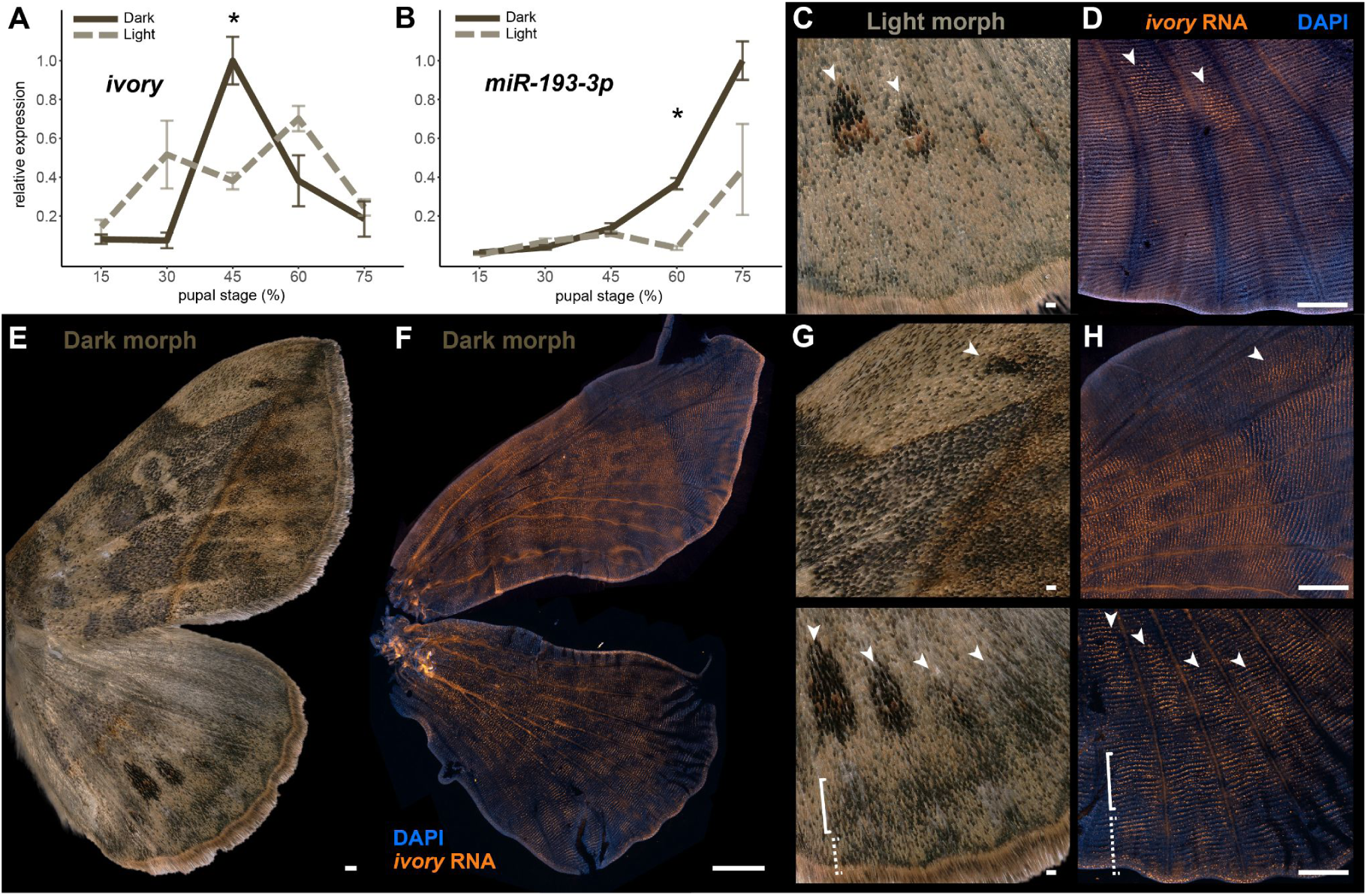
Developmental expression of *ivory:mir-193* in *A. gemmatalis* pupal wings. **(A-B)** RT-qPCR profiling of the *ivory* lncRNA (A) and the mature *miR-193-3p* transcript (B) across 5 stages of pupal development (%) in Light and Dark selected lines. Normalized mean values ± SEM are shown for 4-10 replicates per stage and condition. Asterisk: *p* < 0.05 (Mann-Whitney U-test). **(C-H)**. HCR profiling of the *ivory* lncRNA (orange) in pupal wings at the 45% stage (D, F, H) prefigures the position of melanic patterns in adult wings (C, E, G). C-D features moths from the Light morph selected line, while E-H features individuals from the Dark selected line. Insets, dorsal hindwing spot region: Light morphs (C,D) and Dark morph (G-H, bottom); anterior dorsal forewing: Dark morphs (G-H, top). Arrowheads: melanic spots; brackets: marginal and sub-marginal melanic patterns of the hindwing. Blue: nuclear counterstaining with DAPI. Scale bars: C-D, G-H = 200 μm; E-F = 500 μm.

Given that *ivory* expression diverged most strongly between morphs at 45% of pupal development, we next focused on this stage to map the spatial distribution of *ivory* transcripts across *A. gemmatalis* wings using Hybridization Chain Reaction (HCR) for *in situ* RNA detection^47^. Wings from Light morphs showed strong expression in presumptive melanic spots of the hindwing (**Fig. 4C-D**). At this stage, the diffuse *ivory* HCR signal expression is difficult to discern from background in the other areas of the wings in these Light morphs. Thus, our HCR data likely capture the strongest spatial domains of *ivory* expression, but do not exclude the possibility of lower, broader, or more developmentally transient expression in additional melanic regions. Importantly, we found strong, detectable expression of *ivory* in Dark moth wings that prefigured the position of melanic patterns (**Fig. 4E-H**). These data reveal how *ivory:mir193* is precisely regulated in association to melanic patterns and with distinct levels of melanism between morphs.

### The *ivory:mir-193* transcript is required for melanic scale specification

Next, we used CRISPR-Cas9 microinjections in early embryos to generate indels via non-homologous end joining at the *miR-193* leading strand (*mir-193-3p*), as well as at the *ivory* conserved Transcription Start Site (TSS). Both experiments resulted in G_0_ mosaic knock-out (mKO) adults with strong reductions of melanic pigmentation (**Fig. 5A-D, Table S2**). G_0_ mKO clones for *mir-193* displayed a complete loss of melanic pigmentation on both wing surfaces, while *ivory* TSS mKOs had strongly reduced, but not completely abolished melanin, suggesting some remnant expression of *miR-193* in these individuals. Importantly, mosaic clones in these G_0_ mutants were lighter than their adjacent wild-type clones (**Fig. 5B-C**). This contrast suggests that all scales express a baseline level of *ivory:mir-193* regardless of both Dark and Light morphs, underlying the background beige-grey coloration of the wing. In addition, the sharp boundaries between clones denote the cell non-autonomous activity of *ivory:mir-193* without cell signaling effects to neighbouring cells^48^.

**Figure 5.**
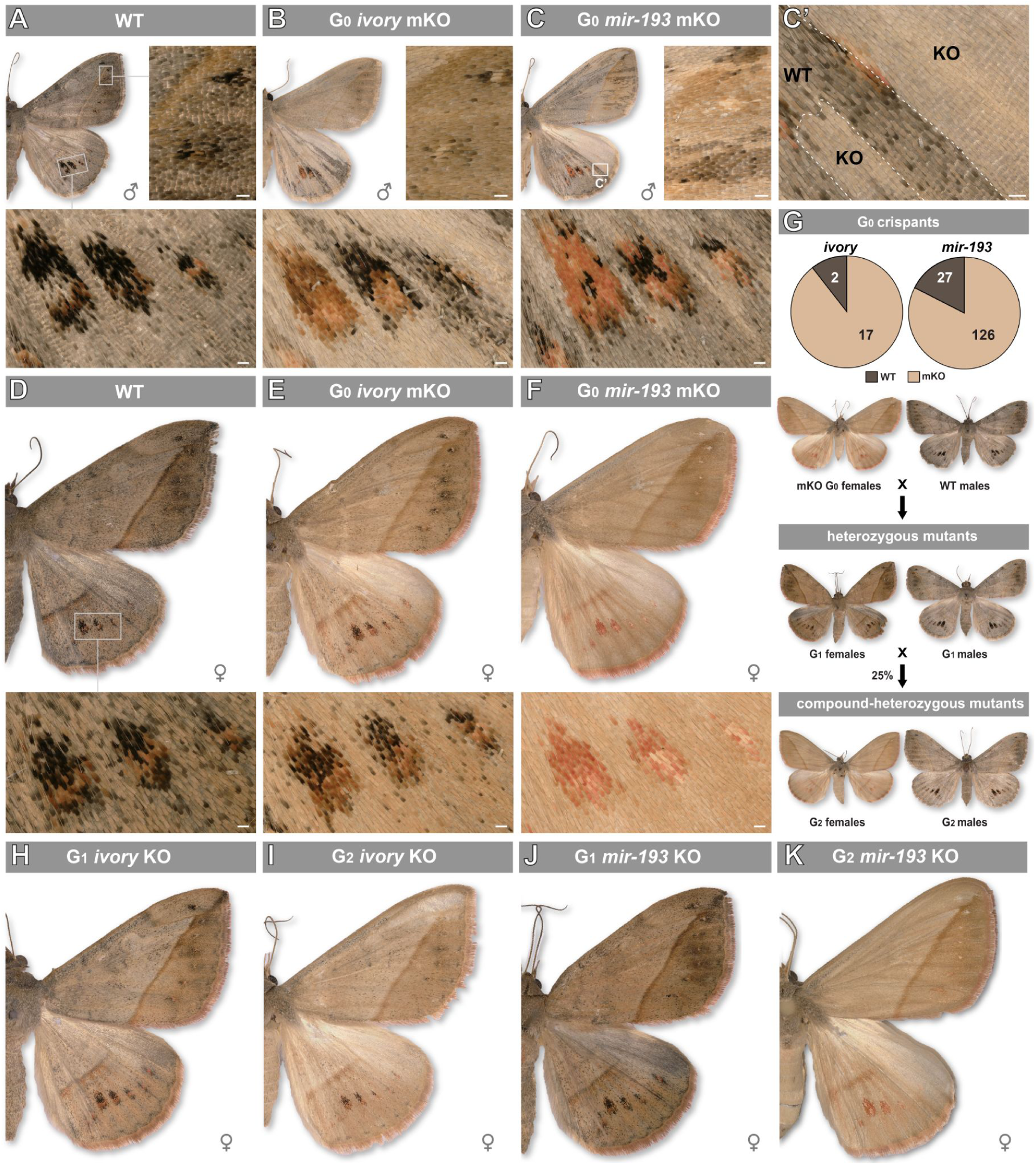
Phenotypic effects of *ivory:mir-193* CRISPR KOs in *A. gemmatalis*. **(A-F)** Dorsal views of wild-type (A, D), G_0_ *ivory* mKO (B, E), and G_0_ *mir-193* mKO moths. Males are shown in the top row, including two individuals with clonal mosaic effects (B-C), with insets featuring magnified views of the forewing distal region (between the M_1_-M_3_ veins) and of the three most prominent hindwing spots. C’: magnified view of the region marked in panel C, with dotted lines marking crispant clone boundaries. Females are shown on the bottom row, including two individuals with crispant clones extending across all wing surfaces. **(G)** Summary of crispant phenotype penetrance obtained in the *ivory* and *mir-193* injection experiments, and subsequent crossing schemes over two generations, resulting in the generation of G_1_ heterozygotes, and G_2_ mutants featuring about 25% compound-heterozygotes. Mutants from the *mir-193* KO experiments are shown here. G_0_ *ivory* mKO mutants were also in-crossed to generate a mixed G_1_ offspring of heterozygous mutants. **(H-K)** Dorsal views of G_1_ and G_2_ *ivory* KO (B, E), and G_1_ and G_2_ *mir-193* female moths, representative of the strongest phenotypic effects obtained at each generation. Scale bars: A-F = 200 μm.

We then out-crossed G_0_ mKO females with WT males, which produced G_1_ heterozygous individuals carrying CRISPR induced deletions for both *mir-193* and *ivory* TSS (**Fig. 5G, Fig. S14-S15**). Heterozygous G_1_ individuals carrying *mir-193* and *ivory* deletions both displayed a reduction in melanic scales as compared to WT individuals, but not a complete loss, suggesting partial co-dominance in expression from both alleles (**Fig. 5H,J**). These heterozygous G_1_ individuals were in-crossed to generate G_2_ compound heterozygous moths, displaying a complete and almost complete loss of melanic scales in *mir-193* and *ivory* mutant lines respectively (**Fig. 5I,K**).

## Discussion

### *ivory:mir-193* is a major-effect locus for camouflage tuning in *A. gemmatalis*

*A. gemmatalis* exhibits striking variation in cryptic wing phenotypes across its range, including differences in hue, overall darkness, and fine-scale textural patterns. Here, we focused on one prominent axis of variation: the amount of dark scale melanin segregating through an outbred lab stock. To dissect the genetic basis for this melanic variation, we used Pool-Seq mapping, long-read phased assembly comparisons, developmental expression assays, and CRISPR-Cas9 perturbations, all of which point to the *ivory:mir-193* locus as the major-effect determinant of melanic camouflage. This species joins a growing list of Lepidoptera where the *ivory:mir-193* hotspot has been linked to adaptive melanism^17,18,23–30,38,41^, and underscores the prevalence of genetic parallelism in driving melanic polymorphisms in butterflies and moths.

Our data suggests that *cis*-regulatory differences upstream of the *ivory* promoter, modulate transcription of a lnc-pri-miRNA precursor, enabling spatial and quantitative production of *mir-193*, which in turn is required for the differentiation of dark scales. This model is supported by three key observations: First, melanic variation maps to a broad interval extending 5’ of *ivory:mir-193*,consistent with segregating *cis*-regulatory modules controlling differences in its spatial recruitmentbetween Light and Dark morphs. Second, *ivory:mir-193* expression is highest in the Dark morphs, and is spatially restricted to melanic patterns in developing wings. Finally, targeted disruption of either the conserved *ivory* TSS or the *mir-193* hairpin region leads to a loss of melanic scales, demonstrating the necessity of the mature *miR-193* for the differentiation of melanic scales. Together, these data argue that *ivory* and *mir-193* operate as a single functional unit, where the lncRNA primarily serves as the transcriptional substrate that is required for *miR-193* production in pattern-specific domains^38^.

This supports a model where *cis*-regulatory differences between Light/Dark alleles spatially regulate *ivory:mir-193* transcription in scale-building cells. These effects cause higher expression levels in the patterns that characterize the Dark morphs, relative to Light morphs where high expression was only observed in the submarginal spots of the dorsal hindwing. While the presence of an allelic series (haplotypes A, B, and C) added statistical noise to the Pool-GWAS, the perfect association of a non-recombining haplotype C with Light phenotypes (**Fig. 2 and Fig. S11**) suggests that the 230-kb haplotype block situated at the 5’ end of *ivory-mir193* is the locus hosting this *cis-*regulatory variation. In future experiments, high-resolution genetic mapping efforts that leverage additional *A. gemmatalis* morphs (**Fig. 1B**), as well as the profiling of open-chromatin and transcription factor occupancy, could decipher how *ivory:mir-193* is spatially regulated to drive specific color morphs.

### Hyperdivergent haplotypes suggest long-term balancing selection on melanism

The cryptic color patterns of *A. gemmatalis* imply a role in camouflage against bird predation^56,57^. Other variable moths showed both reduced predation risk^50,51^ and inter-annual abundance fluctuations compared to monomorphic species^59–61^, suggesting that color polymorphisms are ecologically relevant features, rather than mere byproducts of mutation and drift. We speculate that polymorphic camouflage in moths can be maintained by two forms of balancing selection: apostatic selection due to predator learning, and spatiotemporally varying selection due to habitat heterogeneity. Apostatic selection involves the bias of bird predation towards the most frequent morphs, and can sustain balanced polymorphism through negative frequency-dependent effects^55,56^. At the same time, crypsis in moths is also shaped by the complexity of light environments and visual backgrounds encountered across their range^30,49,57–59^. This spatial heterogeneity likely includes a temporal dimension, as seasonal shifts in vegetation reshape the resting substrate against which avian predators search for prey^60^, a dynamic that may subject the melanic locus to fluctuating selection^61^. Rather than favoring a single optimum, an interplay of frequency-dependent predation and spatiotemporally shifting adaptive landscapes may thus impose strong balancing selection on the *ivoy:mir193* locus, thereby promoting the persistence of multiple morphs in *A. gemmatalis*.

The melanic locus showed the highest level of nucleotide diversity (π) across the genome (**Fig. 2B and Figs. S8-S9)**, suggesting a deep coalescence time between Dark and Light alleles. Long-term balancing selection, by maintaining high levels of genetic diversity within populations over a prolonged period^62,63^, could explain such blocks of divergence. Indeed, long term balancing selection was proposed to underlie the formation and persistence of hyperdivergent haplotypes^64,65^, other examples of color polymorphism^66–70^, and in a strikingly parallel case, camouflage variation linked to the *ivory-mir-193* locus itself in *Kallima* butterflies^18^. Whereas multiple *ivory:mir-193* inversions have introgressed between hybridizing lineages of *Heliconius* butterflies^71–75^*, A. gemmatalis* is the only New World representative of its genus, making introgression a less likely explanation. Rather, we speculate that the hyperdivergent haplotypes at *ivory:mir-193* reflect prolonged balancing selection acting within a single species lineage, preserving deeply divergent Dark and Light alleles over an extended evolutionary time.

### Suppressed recombination maintains polymorphism

Large-effect color patterning loci in insects often include inversions that can reduce recombination and maintain complex *cis*-regulatory haplotypes^18,46,76–79^. Several cases of inversion polymorphism have now been described at the *ivory:mir-193* locus, including the complex evolution of the *Heliconius numata* P supergene inversion^71–73^, a convergent inversion in the *Heliconius erato* clade^73–75^, a ∼1.0 Mb inversion tightly associated with color-pattern polymorphism in the moth *Chetone histrio*^27^, and multiple independent structural haplotypes underlying leaf-mimicry strategies in *Kallima inachus*^18^. *A. gemmatalis* further illustrates this principle: immediately upstream of *ivory:mir-193*, we found complex structural variation between the Dark haplotype A and Light haplotype C, including a double-inversion where the haplotype B lineage acted as an intermediate step (**Fig. 3**). Extreme pairwise nucleotide divergence accompanies this 122-kb wide structural variation, as well the co-linear portion of the haplotype block, where 500-bp alignment windows showed only 75-95% pairwise identity (**Fig. 2D**). These levels of divergence are expected to impair homologous pairing and crossover formation, indicating that both the double-inverted and co-linear regions act as effective barriers to recombination. This effect is most visible in haplotype C, which based on Pool-Seq genotyping, showed complete integrity and absence of recombination with A-B haplotypes beyond the SV region (**Fig. S11**).

In summary we found that large cis-regulatory haplotypes of *ivory:mir-193* differentiate Dark and Light morphs of *A. gemmatalis*, and that their genomic integrity is enabled by suppression of recombination. Consistent with previous cases of hyperdivergent loci^46,64,67,68^, this intrinsic feature likely acts in combination with balancing selection to explain the maintenance of polymorphic camouflage in *A. gemmatalis*.

### Limitations of the study

Our genetic mapping was performed using a laboratory-maintained outbred stock. We identified a main haplotype group underlying the Light-recessive phenotype in this population but unexpectedly, two haplotype groups underlied Dark phenotypes. It is worth noting there is visible variation within each morph in our population (*e.g.* **Fig. 1D**). We suspect that combinations of Dark-associated haplotypes A and B may underlie this variability, an hypothesis that will be testable by generating additional crosses and phased genomes. More broadly, further genetic studies of additional morphs found in nature will also be required to assess if other alleles of this locus contribute to the outstanding level of polymorphism observed in this species, and to estimate their times of divergence.

In addition, the ecological function of the Light and Dark morphs remains to be formally tested. Because these phenotypes are presumed to contribute to crypsis, it will be important to evaluate their camouflage performance across relevant visual backgrounds and habitats. Such experiments could clarify whether the polymorphism is maintained by background-dependent selection, habitat heterogeneity, predator-driven apostatic selection, or other ecological mechanisms.

## Resource Availability

### Lead Contact

Further information and requests for resources and reagents should be directed to, and will be fulfilled by, the lead contact, Luca Livraghi.

### Materials Availability

This study did not generate new materials.

### Data and Code Availability

- The *ilAntGemm2* NCBI RefSeq assemblies of the *A. gemmatalis* genome are available on NCBI Datasets under accession numbers GCF_050436995.1 (primary assembly, annotated), and GCA_050436975.1 (alternate haplotype).
- Small RNA-seq reads are available on the NCBI SRA repository under BioProject PRJNA1165401.
- Pool-Seq reads of Dark vs. Light phenotype females are available under BioProject PRJNA1338508.
- PacBio HiFI reads of 8 F_2_ individuals are available under BioProject PRJNA1450809, and their phased assemblies and corresponding gene and TE annotations are available on the Open Science Framework repository^80^.

## Acknowledgements

We thank the USDA-ARS Ag100Pest Initiative for genome sequencing and assembly; Coralis Acervedo, Rachel Canalicchio, and personnel from the Harlan Greenhouse for growing host plants; Lucas Micheels and John Lill for discussion on moth melanism and assistance pinning specimens; Chad Finkenbinder at Benzon Research for rearing advice; and Tyler Simmonds and Renee Corpuz for the preparation of PacBio and HiC sequencing libraries. The annotation of the *A. gemmatalis* genome was graciously carried out by the NCBI Eukaryotic Genome Annotation Pipeline team. This work was funded by the NSF grants IOS-1656553, MCB-2217156, and a GWU University Facilitating Funds award to AM; by the USDA-ARS Ag100Pest Initiative from the USDA’s Agricultural Research Service; and by USDA-ARS project numbers 2040-30400-003-000D, 0201-88888-003-000D, and 0201-88888-002-000D.

## Author Contributions

Conceptualization, Writing – original draft: L.L., J.J.H. and A.M. Investigation: L.L., J.L.C., M.B., A.T.C., K.K., A.M. Formal analysis: L.L., J.J.H., J.D.A., S.B.S. Funding acquisition and Supervision: S.M.G. and A.M.

## STAR★Methods

## KEY RESOURCES TABLE

Attached separately

## EXPERIMENTAL MODEL AND STUDY PARTICIPANT DETAILS

Larvae of *A. gemmatalis* were commercially obtained from Benzon Research, and originate from a stock established at the USDA Southern Insect Management Research Unit in Stoneville, Mississippi. Benzon Research established an outbred colony of this strain in 2005 and there has been no addition of wild-caught specimens. This stock has been continuously reared as an outbred population with cohorts of around 1,500 reproductive individuals each week (Chad Finkenbinder, personal communication).

All life stages were maintained in a growth chamber at 28°C and 60-70% relative humidity under a 14:10 h light:dark photoperiod. Pupae are separated by sex after examination of gonadal characters on their ventral abdomen, and left to emerge as adults before mixing of 20-40 adults in a 32 ✕ 22 ✕ 15 cm reptile feeding box, supplemented with a 1.6% (m/v) Gatorade powder solution. Males older than 3 days mate and produce viable offspring, while letting adults of both sexes emerging together can produce inviable eggs. Cuttings of soybean (*Glycine max*) and hyacinth bean (*Lablab purpureus*) were provided for oviposition^81^, with their stems inserted in floral tubes filled with a hydrogel to maintain hydration (we recommend using the content of gel ice packs used for shipping for this purpose). Egg laying occurs preferentially within the first hours of the scotophase. Eggs are collected from leaves by hand over a glass dish, washed with 5% Benzalkonium Chloride for 1 min, rinsed with water and dried before microinjection or rearing. For rearing, embryos were kept in a container with saturating humidity for 48 h, before addition of *L. purpureus* cuttings for hatching (at 28°C, about 72-80 h after egg laying, AEL), and feeding. Larvae were transferred at their second or third instar to single 37 mL cups Multiple Species Diet (Southland Inc) prepared with approximately 90% of the manufacturer-recommended water volume (*e.g.* 850 mL instead of 930 mL) to reduce humidity within feeding containers, and with lids pierced with forceps for aeration. Alternatively, embryos were left to hatch on hydroponic trays of 7–21 d old vetch sprouts^82^, placed inside a collapsible mesh cage, before transfer to cups at the fourth instar. Pupa formation occurs at the end of the fifth instar stage. Pupal development takes on average 171 h in females and 193 h in males at 28°C.

## METHOD DETAILS

### Hybrid crosses

Adult individuals obtained from the outbred commercial stock (Benzon Research) were separated into breeding pools of 10-20 Light and Dark individuals for two generations, resulting in two stocks with pure-bred phenotypes. A Dark female was crossed to a Light male to generate F_1_ offspring, from which sib-sib pairs were used to generate two F_2_ families (brood 3D, N = 25; brood 3F, N = 53). Similarly, a Light female was crossed to a Dark male to generate F_1_ offspring, from which sib-sib pairs were used to generate two F_2_ families (brood 6A, N = 32; brood 6C, N= 11).

### Reference genome sequencing and assembly

High-molecular weight DNA was extracted from an *Anticarsia gemmatalis* female pupa, following two generations of inbreeding between Dark morphs, using a Qiagen Genomic-tip 100/g. High-molecular weight DNA resulting from the extraction was qualified and quantified using the Genomic DNA 165-kb Kit on an Agilent Femto Pulse system and Qubit Broad Range dsDNA kit on a Denovix DS-11 along with UV spectrophotometric readings for purity. Approximately 500 ng of DNA was subjected to shearing on a Diagenode Megaruptor 3, targeting 15 - 20 kb fragments and prepared for PacBio HiFi sequencing using a SMRTBell Express Template Prep 3.0 kit. The resulting library was sequenced on a PacBio Revio system.

For Hi-C sequencing, a pool of 5 Dark morph, female pupal heads were snap-frozen in liquid nitrogen and stored at -80°C before preparation and sequencing. The HiC library was prepared following a standard protocol^83^ using the restriction enzymes DdeI and DpnII, and the resulting proximity ligation library was prepared for sequencing using the NEBNext Ultra II DNA Library Prep Kit. The final libraries were sequenced on a partial flow cell using the AVITI 2×150 Sequencing Kit Cloudbreak FS High Output kit on the Element AVITI System. Following sequencing, raw reads were basecalled using *bases2fastq* v.1.3.0.

Sequencing reads from one PacBio Revio SMRTCell were filtered for adapter contaminants using NCBI FCS-Adaptor and HiFiAdapterFilt ^84,85^, and assembled into contigs using HiFiASM v. 0.19.3-r572^86^. Duplicate contigs were identified and removed from the assembly using PurgeDups^87^. Sequenced HiC reads were mapped to the duplicate purged contigs using BWA-mem2^88^ and a HiC contact map was created using the YAHS pipeline^89^. Assembled scaffolds were deposited with the National Center for Biotechnology Information (NCBI) under accession GCA_050436995.1. Contigs corresponding to an alternate haplotype assembly of 113.7 Mb were also deposited online (NCBI: GCA_050436975.1). The assembly has a BUSCO v5.8.0 ^90^ completeness score of 99.3% (single-copy 99.1%, duplicated 0.2%) using the *lepidoptera_odb10* reference ortholog dataset (n=5,286). The BUSCO ortholog assignment results were used to assign lepidopteran ancestral linkage groups, termed Merian elements, to each chromosome with *Lep_BUSCO_Painter* v.1.0.0^91^.

### RNA-seq and genome annotation

To assist with the functional annotation of this genome, we generated RNA-seq transcriptomes from embryos, larval heads, body and silk glands, adult heads, testes, and ovaries, and forewing tissues from pupae dissected that 40-80% stages of pupal development in 10% increments. For RNA-seq, dissected tissues were stored in TRI-Reagent and stored at -70°C, before total RNA extraction, cDNA preparation, library preparation, and Illumina NovaSeq sequencing at a target volume of 20M PE150 reads by Genewiz (South Plainfield, NJ). Out of 16 libraries, 13 were prepared from cDNA prepared from oligo-dT reverse transcription; 3 RNA-seq libraries were generated using cDNA libraries generated by random priming, but generated more than 90% of rRNA reads due to the failure of the QIAseq FastSelect Fly Kit to deplete ribosomal RNA in our species. Dissected tissues were stored in TRI-Reagent at -70°C. A functional annotation of the *A. gemmatalis* genome (NCBI RefSeq: GCF_050436995.1) was then generated by the NCBI Eukaryotic Genome Annotation Pipeline initiative^92^.

### Small RNA-seq

For annotation of the mature *miR-193* in the genome, small RNA-seq transcriptomes were generated from 40% developed pupal wings, adult testes, and ovaries. Dissected tissues were stored in TRI-Reagent and stored at -70°C. Total RNA extraction, cDNA generation and barcoding via the Small RNA library Preparation Kit (New England Biolabs), and paired-end Illumina NovaSeq sequencing were outsourced to Genewiz (South Plainfield, NJ), using the small RNA-seq sequencing service with a target yield of 10M paired-end reads per sample. Read quality was assessed with *FastQC v0.12.1*, before adapter and poly-G tail trimming with *Flexbar v3.5.0* using a FASTA file containing the Illumina small RNA-seq-specific adapter sequences and with the option *-ap ON* ^93,94^. *FastQC* was used post-trimming to validate adapter sequences and polyG tails had been removed. The trimmed reads were aligned to the genome assembly (GCA_050436995.1) using *STAR v2.7.11b* ^95^. The alignment BAM files were then converted to SAM format using *Samtools v1.22* and then converted to an ARF file format using the bwa_sam_converter Perl script from *miRDeep2* ^96,97^. All available lepidopteran miRNA hairpin and mature sequences were downloaded in FASTA format from miRBase in November 2025^98^. To identify, annotate, and fold miRNA precursors in the genome, *miRDeep2* was run with a fasta file specifying the *miR-193* star sequence that was manually identified using the BAM files previously generated in IGV and in comparison with other known lepidopteran *miR-193* star sequences and the miRbase lepidopteran miRNA sequences ^38,98^. Geneious Prime was used to visualize the results of the STAR alignment and *miRDeep2* annotations in the 3’ region of the *ivory* lncRNA.

### Pool-GWAS mapping of the major-effect melanic locus

To identify genomic regions associated with melanic scales, we performed a pooled whole-genome association analysis (Pool-Seq) contrasting Dark and Light morphs (25 Dark females *vs.* 23 Light females). Genomic DNA from individuals representing each phenotype was extracted using a Zymo Quick DNA Miniprep Plus kit and combined into two pools (one per phenotype), with an equal DNA concentration from each individual. Library preparation and sequencing of each pool was performed by Genewiz to an average normalised depth of 88X, calculated using *mosdepth*.^45^ Reads were subsequently aligned to the *A. gemmatalis* reference genome using *BWA-MEM*^99^, and quality and coverage filtering performed with *samtools*^96^. Allele frequency differences between the Dark and Light pools were calculated using the *PoPoolation2* pipeline.^100^ Briefly, allele frequencies were called with *samtools mpileup* and quantified using the *PoPoolation2* implementation of Fisher’s exact test, applied per site across the genome, and association strength was summarized as –log₁₀(P) for each site. Because our Pool-Seq design consisted of a single two-pool comparison, we did not impose an empirical genome-wide significance cutoff on windowed association values, as site- and window-level *P*-values can be strongly influenced by coverage heterogeneity and other sources of background inflation. Instead, we used per site *P*-values in order to retain local structure in the association profile and defined the GWA interval from the contiguous peak shape in combination with independent differentiation metrics, including F_ST_ and D_XY._ Pool-GWAS mapping was repeated using the same pipeline against our assembled Light Reference genome to rule out any assembly specific associations.

### Calculation of population genetics statistics

Pool-seq reads from Dark and Light morph pools were aligned to the Dark reference genome and processed using the same filtered allele-count dataset used for Pool-seq GWAS. Window-based estimates of genetic differentiation and divergence, including F_ST_, D_XY_, and nucleotide diversity π, were calculated using *Grendalf*, in 10-kb windows with a 1-kb slide. Calculations were performed on genome-wide distributions and compared between windows overlapping the chromosome 8 GWA interval containing *ivory:mir-193* and the rest of the genome.

### PacBio HiFi long read-read resequencing and phased haplotype assemblies

High-molecular-weight DNA was extracted from 8 F_2_ adult females derived from two independent crosses (2 Dark and 2 Light per brood, from broods 3F and 6A) using a Qiagen Genomic-tip 100/G kit. Libraries were prepared as described for the reference sample and multiplexed for sequencing on a single PacBio Revio SMRT Cell. Sequencing produced between 30.3× and 48.9× HiFi coverage per individual, with a mean coverage of 40.6×. HiFi reads from each of the 8 F_2_ individuals, as well as the raw reads from the Dark reference genome individual, were assembled independently using *Hifiasm*^86^ in default HiFi parameters as follows. Briefly, *PacBio HiFi* BAM files were converted to FASTQ, filtered for residual adapter contamination, and assembled without the *--primary* or *-l0* flags, since our goal was to preserve local haplotype structure rather than collapse the assembly to a single pseudohaplotype representation. We retained the haplotype-specific assembly outputs together with haplotype-resolved unitig graphs for downstream analysis. For each individual, GFA segment records were converted to FASTA, and *cortex*-containing haplotype contigs were identified by BLAST for structural comparison. This yielded two phased haplotypes per individual, for a total of 16 haplotypes across the 8 genomes.

For assembly of the Light reference genome, the individual with the highest PacBio HiFi yield (female *Light 04*; 48.9× coverage) was selected and assembled *de novo* with *Hifiasm* using default parameters to generate a pseudohaplotype assembly. To enable direct chromosome-level comparison with the Dark reference genome, the resulting contigs were ordered and oriented relative to the Dark chromosome-level assembly using *RagTag*^103^, producing a reference-guided pseudomolecule assembly that retained Dark-reference chromosome naming and order. Gene annotation was then transferred from the Dark reference genome to the scaffolded Light assembly using *Liftoff*^104^.

### Generation of genotype plots and haplotype frequency calculations

To visualize haplotype structure across the associated interval, phased *PacBio HiFi* haplogenomes from resequenced individuals were aligned independently to the Dark reference genome using *minimap2*, with each *Hifiasm*-resolved haplotype FASTA treated as a haploid assembly. Alignments were sorted and indexed with *Samtools*96, and haplotype-level variants were called relative to the Dark reference using *bcftools mpileup* and *bcftools call*96. Calls were filtered to retain sites supported by a single aligned haplotype sequence at each reference coordinate (DP = 1), while sites with no coverage, multiple coverage, or ambiguous calls were treated as missing. Individual haplotype VCFs were normalized, merged into a multi-sample VCF, filtered to retain informative biallelic SNPs, and subset to the chromosome 8 GWA interval. This VCF was used as input for *GenotypePlot*105, with genotypes plotted relative to the Dark reference allele.

To determine the frequencies of the B and C alleles in the Pool-Seq data, we selected all private fixed SNPs in both B and C haplotypes within a pre-defined haplotype block (2.34 Mb-2.57 Mb). We scored the occurrence frequency of each of these SNPs, and used the median frequency of all sites to infer a percentage-occurence of the B and C alleles in each pool.

### Analysis of structural variation at the ivory:mir-193 locus

*Mosdepth*^45^ was used to quantify read depth in each pooled library from the final BAM files. BAMs were indexed with *samtools* and median depth was calculated in 100-bp windows using *mosdepth* (--use-median --by 100), producing BED-format outputs. To compare pools, window median depths were normalized within each library by the chromosome-wide baseline depth estimated from all windows on Chromosome 8 (median of non-zero windows), such that normalized depth values are centered near 1. To test whether depth depletion within the Pool-GWA interval was preferentially associated with non-coding sequence, exon annotations on scaffold NC_134752.1 (chromosome 8) were extracted from the GFF, converted to BED, and merged to define a non-redundant set of exonic and non-exonic sequence (intronic + intergenic). Exon-only and non-exon-only windows were partitioned into those inside versus outside the Pool-GWA interval, and *mosdepth* was used to compute window-level depth distributions.

For comparison of long-read haplotypes, the contig containing the *ivory:mir-193* locus was identified by BLASTing a *cortex* exon against each phased assembly and extracting the associated contigs. Equivalent regions from the dark and light reference genomeswere aligned with *MUMmer4*^105^ and visualized as a dotplot. Pairwise sequence similarity among phased haplotypes across the candidate region was visualized using dotplots generated with *blastn2dotplots*^106^. For each comparison, the extracted haplotype sequence was aligned against the homologous interval from the Dark reference assembly using *BLASTN*, and alignments were plotted with *blastn2dotplots* using a minimum alignment identity threshold of 70% and a minimum alignment length threshold of 500 bp. All haplotypes were oriented and compared relative to the Dark reference coordinate system, allowing direct visualization of collinearity, local inversions, and highly divergent segments among phased assemblies.

### Transposable element quantification and syntenic comparisons of inversions

Repeats were annotated in the Dark reference genome and three representative haplogenomes (A–C) using *EarlGrey* (REF). A species-specific de novo repeat library was first generated from the Dark reference and then combined with the *Dfam* Obtectomera database for annotation of the Dark reference and representative haplogenomes. This combined repeat library was then used in subsequent *EarlGrey* annotation-only runs to annotate the representative haplogenomes. Filtered *EarlGrey* GFF outputs were used for all downstream analyses. Repeat annotations were grouped into broad classes (DNA, LINE, LTR, rolling-circle, SINE, simple repeats, and unclassified), and basepair coverage was calculated as the union of annotated bases within each class. We then quantified repeat composition genome-wide and within the chromosome 8 GWA interval for each representative genome. To test whether the focal interval was repeat-rich, we compared repeat coverage in the GWA interval to genome-wide distributions of matched windows generated with window sizes equal to the corresponding GWA interval and a 50-kb step size. For each haplotype group (A–C), we calculated repeat-covered fraction for each major repeat class and for total repeat content across all windows.

For synteny visualization between haplotypes, anchored intervals surrounding the inversion breakpoints were aligned pairwise with *LASTZ* using gapped, both-strand, chained alignments. AXT alignment output was plotted with *ggplot2* as ribbon-style synteny plots linking homologous coordinates between sequences. Gene models (exon coordinates) and TE annotations from the composite *EarlGrey* GFF files were overlaid on each interval, allowing synteny patterns to be interpreted relative to local gene structure and repeat content.

### Real-time quantitative PCR

RT-qPCR was performed on pupal forewings of Light and Dark line female individuals dissected at 15%, 30%, 45%, 60%, and 75% of pupal development, with 3-5 individuals used as biological replicates for each stage and condition. For accurate staging, pre-pupae were transferred to transparent cups and pupation time was recorded using a TX-164 timelapse camera (Technaxx), and dissected at the wanted stage based on a total pupal development period of 171 h. Individual forewings were dissected in ice-cold Phosphate Buffer Saline (PBS 1X), transferred to 800 μL TRI-Reagent for RNA stabilization, and stored at -70°C. Total RNA was isolated using the Direct-zol RNA Miniprep kit (Zymo Research) and stored at -70°C. RNA concentrations were determined via NanoDrop. From each sample, 1 μg of RNA was converted into cDNA using the PrimeScript RT Reagent Kit with gDNA Eraser (TaKaRa Bio). The manufacturer reverse transcription primers (random hexamers and poly-dT) were used to generate cDNA libraries for quantification of the *ivory* lncRNA. Stem-loop primers were used for the reverse transcription of the mature *miR-193-3p* miRNA^38,107^. RT-qPCR was performed using Luna Universal qPCR Master Mix (New England Biolabs) following manufacturer recommendations). To quantify *ivory* and *ebony* expression, gene-specific primers were used for amplification, and normalized to *Rsp18* expression, using a thermocycler program set at 40 cycles of 95°C for 15 s and 60°C for 30 s. For *miR-193* expression, *miR-193-3p* and stem-loop primers were used for amplification, with the *U6* RNA for normalization^38^, using a thermocycler program set with 40 cycles of 95°C for 15 s and 55°C for 30 s. Two technical replicates were performed for each experiment, on at least 4 biological replicates per stage and condition.

### Hybridization Chain Reaction (HCR) in situ hybridization

To ensure accurate developmental timing for dissections, total pupation time was recorded from onset of pupa formation to eclosion using a TX-164 timelapse camera (Technaxx). Total development time was used to calculate pupal % development. Ten HCR probes were designed against the *ivory* coding exon1 (**Table S3**), using the *insitu_probe_generator tool*^108^, permitting a GC content in the 35-75% range, and allowing limiting runs of poly-GC and poly-AT to a maximum of 3 bp. HCR^47^ was performed as previously described with minor modifications^109^. Briefly, pupal wings were dissected from the pupal case in cold 1X PBS, transferred to a fixative solution (750 μL PBS 2 mM EGTA, 250 μL 37% formaldehyde) containing 9.25% formaldehyde at room temperature for 30 min and washed four times in PBS containing 0.01% Tween20 (PBT). Pupal wings were then permeabilized in 1 μg/μL of ProteinaseK diluted in PBT solution for 5 min at 37°C. Samples were then washed with a stop solution containing PBT and 2 mg/mL glycine and followed by two additional PBT washes. After transferring wings to a post-fix solution (850 μL PBT, 150 μL 37% formaldehyde) containing 5.55% formaldehyde for 20 min, wings were washed four times with PBT before following the rest of the standard protocol^110^. Samples were then washed in PT, incubated in 50% glycerol with with 1 μg/mL DAPI (4′,6-diamidino-2-phenylindole), mounted on glass slides with 70% glycerol or SlowFade Gold mountant under a #1.5 thickness coverslip, and sealed with nail varnish.

### CRISPR mosaic knock-outs and generation of mutant lines

Embryo microinjections followed published procedures^82,111,112^ using CRISPR injection mixes with 2:1 mass ratios of Cas9-2xNLS and synthetic sgRNAs, targeting the *ivory promoter* and the *mir-193* hairpin region (**Table S4**). Synthetic sgRNAs were provided by Synthego and resuspended in Low TE buffer (10 mM Tris-HCl, 0.1mM EDTA, pH. 8.0) in 2.5 μL aliquots (500 ng/μL) and stored at - 70°C. Cas9-2xNLS recombinant protein (QB3/Macrolabs, UC Berkeley) was resuspended at 1,000 ng/μL in *Bombyx* 2X injection buffer (0.15 mM NaH₂PO₄, 0.85 mM Na₂HPO₄, and 10 mM KCl, pH 7.2–7.4) containing 0.05% cell-culture grade Phenol Red, and stored at -70°C as 2.5 μL aliquots. Equal volumes of aliquoted Cas9 and sgRNA aliquots were mixed to a final concentration of 500:250 ng/μL before microinjection.

Injections were performed before blastoderm formation around 4 h, with a majority of injections completed within 2 h after egg laying. Survival and phenotypic penetrance were assessed in larvae, pupae, and adults, and no morphological phenotypes were visible in immatures in this study. Resulting G_0_ mosaic KOs (mKOs) for both *ivory* and *mir-193* were then crossed to wild-type (WT) moths to establish mutant lines. Specifically, G_0_ mKO females were outcrossed to WT males to generate G_1_ offspring, among which heterozygous individuals were identified based on reduced melanic pigmentation and subsequently confirmed by genotyping (see below). G_1_ individuals exhibiting the strongest and most consistent reduction in melanism were then intercrossed to produce G_2_ progeny, yielding individuals segregating a range of CRISPR-induced deletion alleles at the targeted locus and enabling recovery of compound heterozygous mutant genotypes for phenotypic characterization.

### Genotyping of CRISPR-induced mutations

Total DNA was isolated from leg tissue using the Quick-DNA Miniprep Plus Kit following manufacturer instructions (Zymo Research) and quantified using a Qubit 3.0 fluorometer (Thermo Fisher Scientific) and the AccuGreen Broad Range dsDNA Quantitation Kit (Biotium). Amplicons spanning the *ivory* CRISPR target sites were amplified in 100 μL PCR reactions using primers *ivory-F* (5′-GAACACGACAACCCCACTTC-3′) and *ivory-R* (5′-CCTCTTGGCTGCATCTTGTC-3′) and purified using the DNA Clean and Concentrator kit (Zymo Research). Sequencing was outsourced to Genewiz for Oxford Nanopore long-read sequencing using the EZ-PCR service. Nanopore reads were mapped to the corresponding *ivory* amplicon reference using *minimap2*, and resulting BAM files were inspected for indel alleles by extracting full-length amplicon reads and generating per-allele consensus sequences. Individuals were classified based on the presence of wild-type and/or deletion consensus alleles (heterozygotes: WT + single deletion; compound heterozygotes: two distinct deletion alleles).

For *mir-193* G_1_ and G_2_ mutants, DNA was extracted and quantified as above, and low-pass whole-genome sequencing was outsourced to Genewiz (150 bp paired-end libraries). Reads were mapped to the *A. gemmatalis* reference genome using BWA-MEM. Local sequencing coverage was quantified using *mosdepth*^45^ in fixed windows and normalized by dividing window depth by the mean depth across the corresponding scaffold. To account for population-level structural variation segregating in the laboratory stock, baseline (“wild-type”) coverage expectations were estimated from the Pool-seq dataset and used for comparison to normalized coverage profiles in G_1_ and G_2_ individuals.

### Microscopy and image processing

Confocal imaging stacks of HCR in situ hybridizations were also obtained on an Olympus FV1200 confocal microscope and a Zeiss LSM 800 confocal microscope, each mounted 10✕ objectives. Fluorescent acquisitions were processed in FIJI. Adjustments of contrast limits were applied independently to each fluorescent channel. Bright-field macrophotographs of whole moth specimens and wing patterns were obtained using Keyence VHX-5000 microscope mounted with VH-Z00 and VH-Z100 lenses, using fixed lighting, exposure and gain conditions that preserved color rendering across samples at each magnification level.

## QUANTIFICATION AND STATISTICAL ANALYSIS

### Scoring of melanic variation using wing scans

The dorsal and ventral surface of forewings and hindwings from select female individuals were imaged on an Epson Perfection V600 scanner, at a resolution of 2,400 dpi, alongside a 24-colour correction card and greyscale standards. Wing images were exported as 8-bit RGBA PNG files with transparent backgrounds and analysed using the R package *recolorize2* ^113^. To quantify dorsal forewing darkness, we used a luminance-based metric derived from the CIELAB color space. For each specimen, we extracted all non-transparent wing pixels (alpha > 0) from each image, converted the corresponding sRGB values to CIELAB using grDevices::convertColor, and retained the L* channel as a per-pixel measure of luminance (lower L* = darker). To summarize darkness per specimen, we calculated the proportion of wing pixels with L* below a fixed threshold (L* < 30). Specimens with visibly damaged wings were excluded from this analysis.

### Probabilities of haplotypes A or B escaping sampling in the Light pool

For haplotypes A and B, we observed the allelic frequencies *fD* in the Dark pool of 50 haploid genomes (25 diploids), and no alleles in the Light pool (*fL* = 0) of 46 haploid genomes (23 diploids). Their respective frequency *f* in a combined population preceding pooling is:

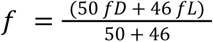

If either of these haplotypes were compatible with the Light phenotype, their probability to escape random sampling from the Light pool would be:

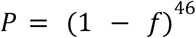

For haplotype A, we observed *fD* = 0.29, leading to:

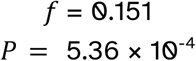

For haplotype B, we observed *fD* = 0.37, leading to:

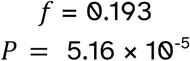

Thus, the probability that haplotypes A or B escaped sampling in the Light pool is low and it follows that A/C and B/C heterozygous states yield a Dark phenotype, *ie.* that A and B both provide Dark-dominant effects. We also know from the reference Dark genome that the A/A homozygous state yields a Dark phenotype, and from the F_2_ genomes, that the C/C homozygous state yields a recessive-Light phenotype. The effect of B/B genotypes was not tested, but the dominance of B over C (B/C = Dark, as inferred from its absence in the Light Pool) suggests that B/B most likely yields Dark phenotypes (B/B = Dark).

### Gene expression profiling

To test for statistical differences in gene expression between Light and Dark morphs, expression values obtained by qPCR were compared independently at each developmental stage using a two-tailed Mann–Whitney *U* test. Statistical tests were performed on ΔCt values, which provide a log-scaled measure of relative transcript abundance. For visualization, ΔCt values were converted to linear expression units (2^−ΔCt^) and plotted as mean ± SEM.

## SUPPLEMENTAL INFORMATION

**Figure S1.**
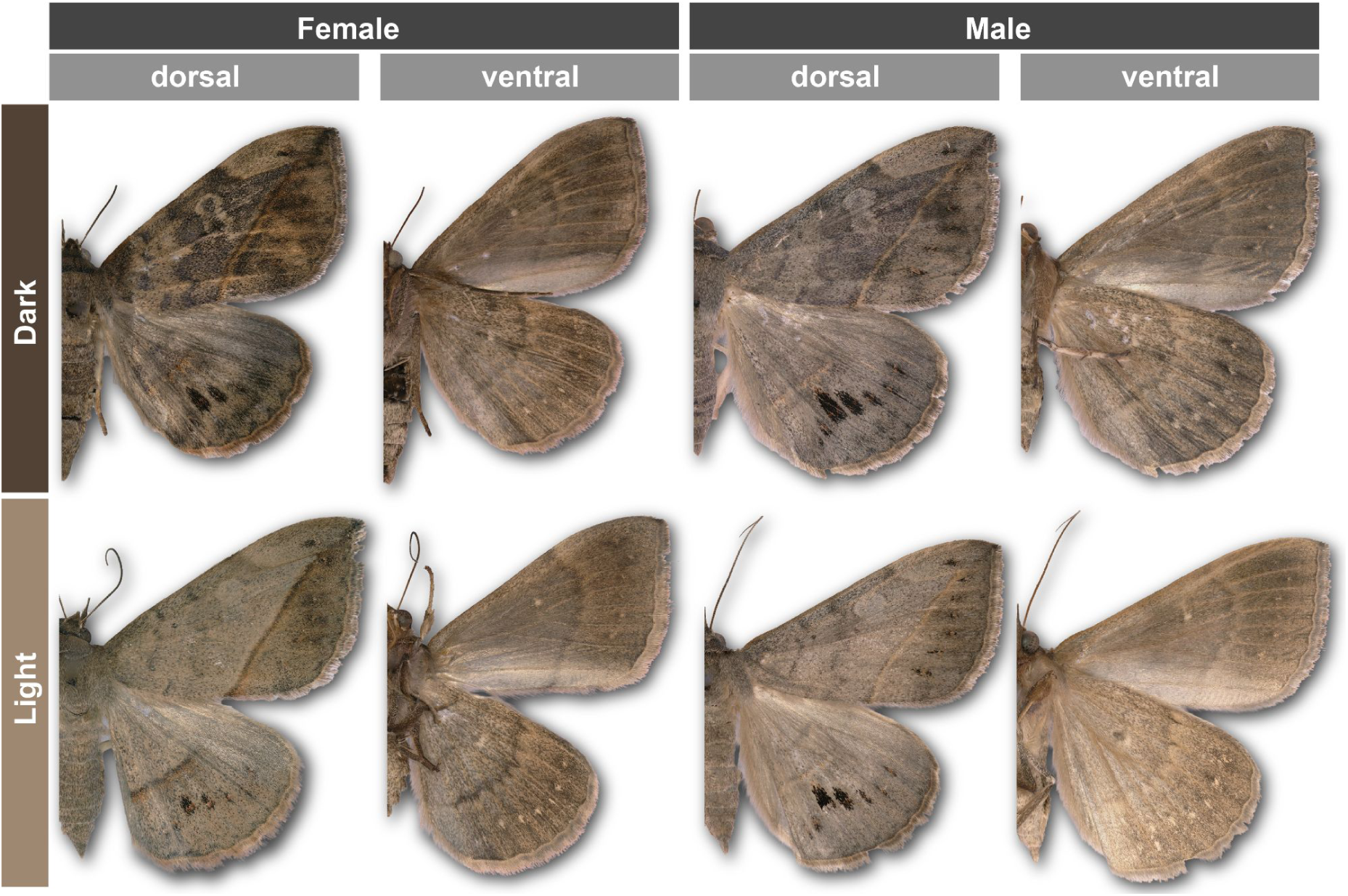
Wing phenotypes in *Anticarsia gemmatalis* segregating in the Stoneville strain. Related to Fig. 1. Representative adults showing the two discrete color morphs (Dark and Light) across sexes and wing surfaces.

**Figure S2.**
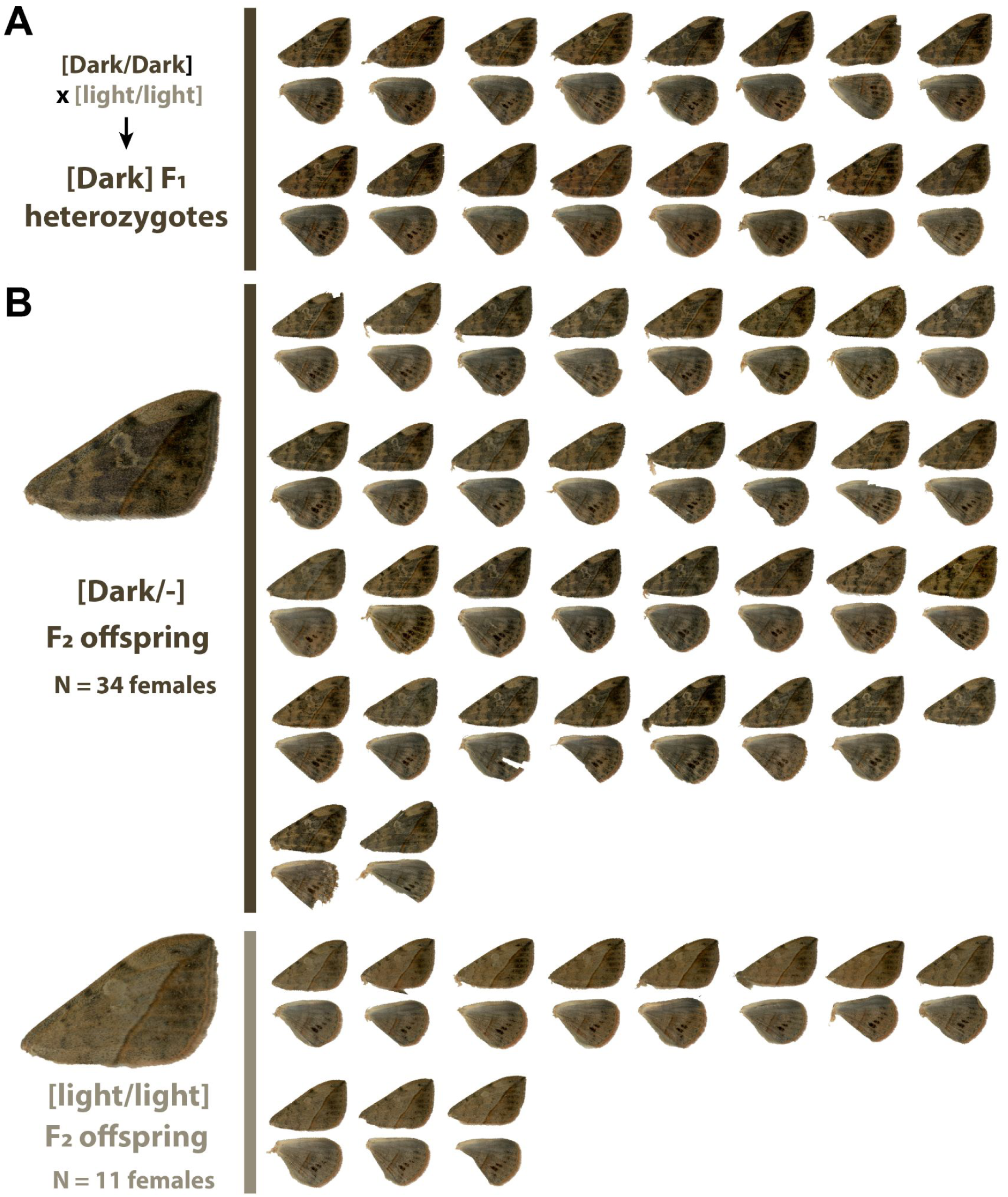
F_1_ and F_2_ hybrid crosses indicate that a large-effect locus controls melanic variation. Related to Fig. 1. F_2_ offspring from a cross between pure-bred Light and Dark morphs resulted in two phenotypic classes. Dorsal wings of female individuals are shown here. **(A)** F_1_ heterozygotes show a Dark phenotype. **(B)** F_2_ offspring resulting from F_1_ sib-matings yield Dark:Light morphs in a 3:1 ratio. Males showed similar effects, and ratios from combined sexes are consistent with a single locus segregating for a dominant Dark allele and a recessive light allele (56 Dark : 18 Light F_2_, Chi-square test for 3:1 ratio, P > 0.89).

**Figure S3.**
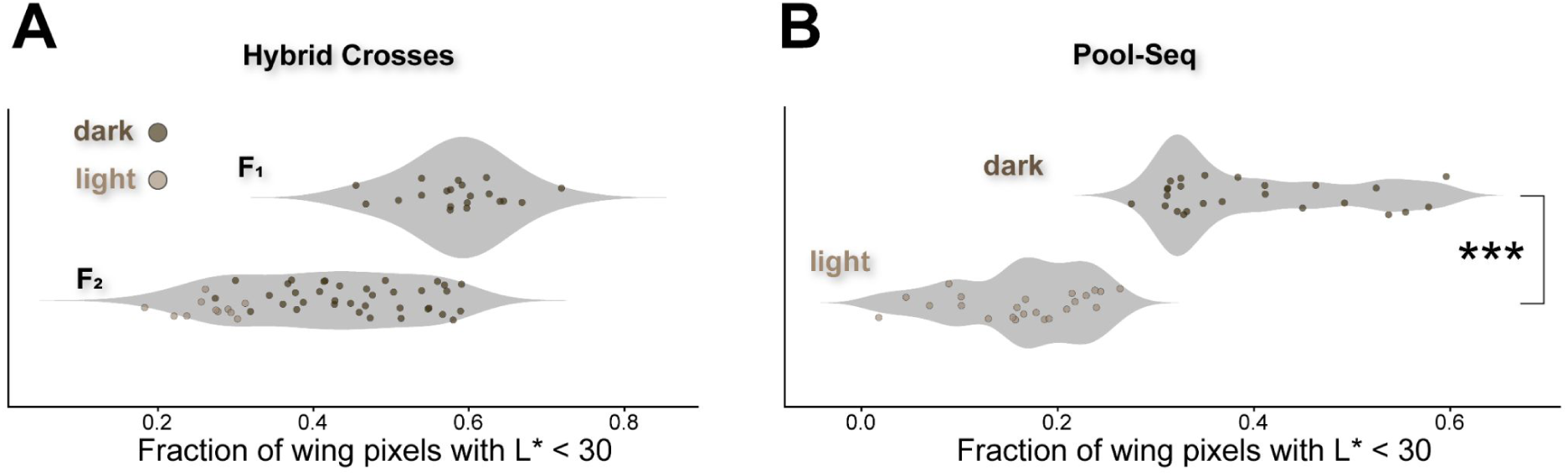
Luminance-based quantification of melanism in hybrid crosses and Pool-seq individuals. Related to Fig. 1. **(A)** Violin plots showing the distribution of per-individual “darkness” scores, quantified as the fraction of dorsal forewing pixels with CIELAB luminance L* < 30. Offspring from a Dark female × Light male cross (F₁; all visually scored as dark) cluster at higher darkness values, consistent with dominant inheritance of the dark phenotype. Sib–sib matings among F₁ individuals produce F₂ offspring spanning the full phenotypic range, including individuals visually scored as Dark or Light morphs. Points represent individuals, with filling colors indicating *a priori* visual phenotype assignments (Dark morphs: brown; Light morphs: cream). The two grandparents and four F_1_ parents used for generating the two F_2_ crosses were not quantified due to wing wear. **(B)** The same metric applied to the individually imaged specimens used for Pool-seq shows clear separation between individuals visually assigned to the Dark and Light morph categories. Group differences were assessed with a Wilcoxon rank-sum test (***, *P* = 3.16 × 10⁻⁹).

**Figure S4.**
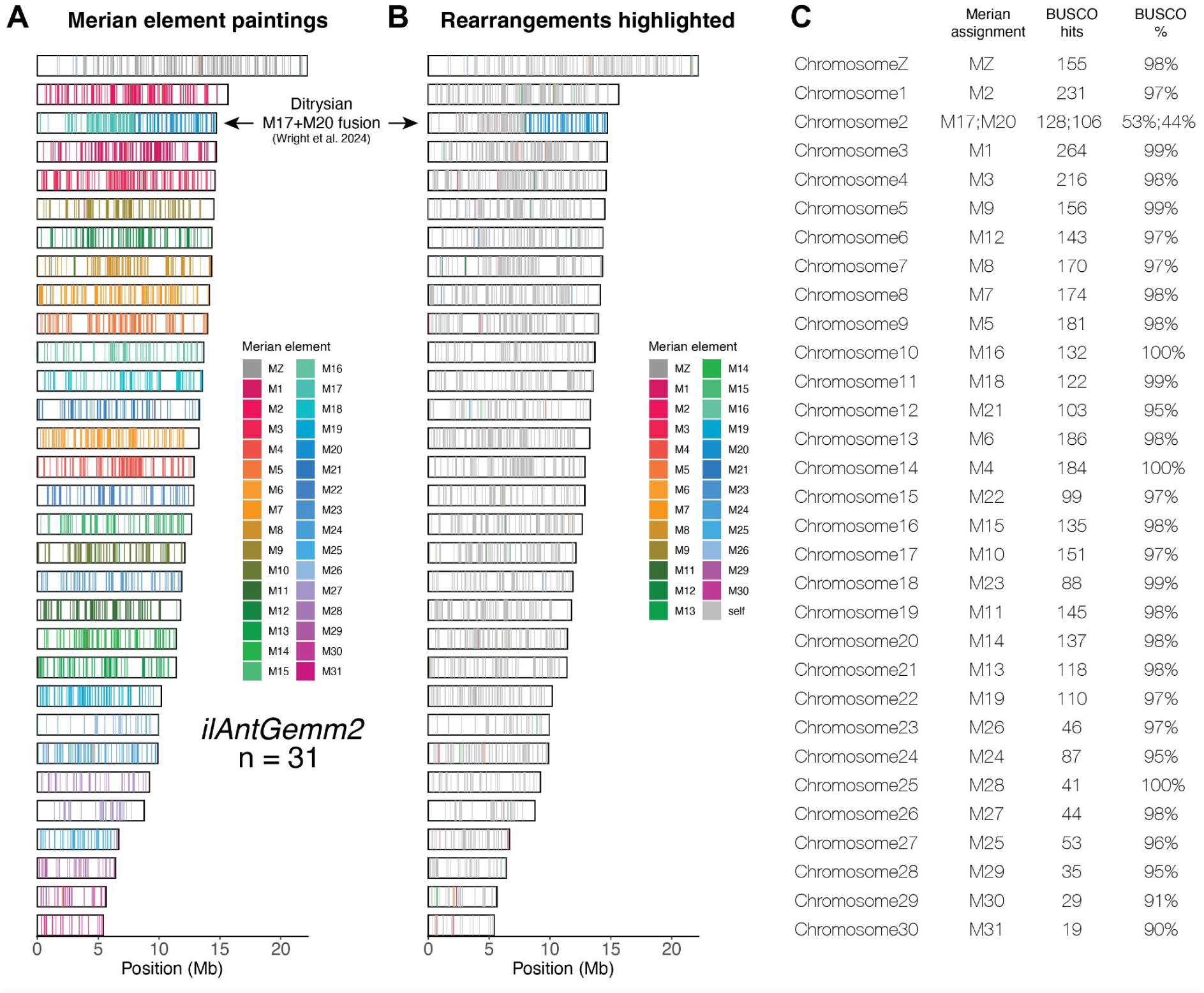
A chromosome-level annotated genome for *A. gemmatalis* moths. Related to Fig. 1. The *ilAntGemm2* primary haploid assembly of *A. gemmatalis* (NCBI: GCA_050436995.1) includes complete chromosome scaffolds and is functionally annotated with 15,773 predicted genes supported by RNA-seq evidence. This resource is the first NCBI RefSeq genome for Erebidae, a large moth family of about 35,000 described species. **(A-B)** Merian elements mapped across chromosomes in the *ilAntGemm2* assembly. N = 31 scaffolds corresponding to 30 autosomes and the Z chromosome are drawn to scale. BUSCO paintings with orthologue positions are shown as coloured bars, each shaded according to its corresponding Merian element (**A**), or with discrepancies with the ancestral lepidopteran linkage groups highlighted (**B**). The M17-M20 fusion is ancestral to Ditrysia, a lineage that encompasses most of lepidopteran diversity ^19,91^. **(C)** Merian assignments for each *A. gemmatalis* chromosome scaffold.

**Figure S5.**
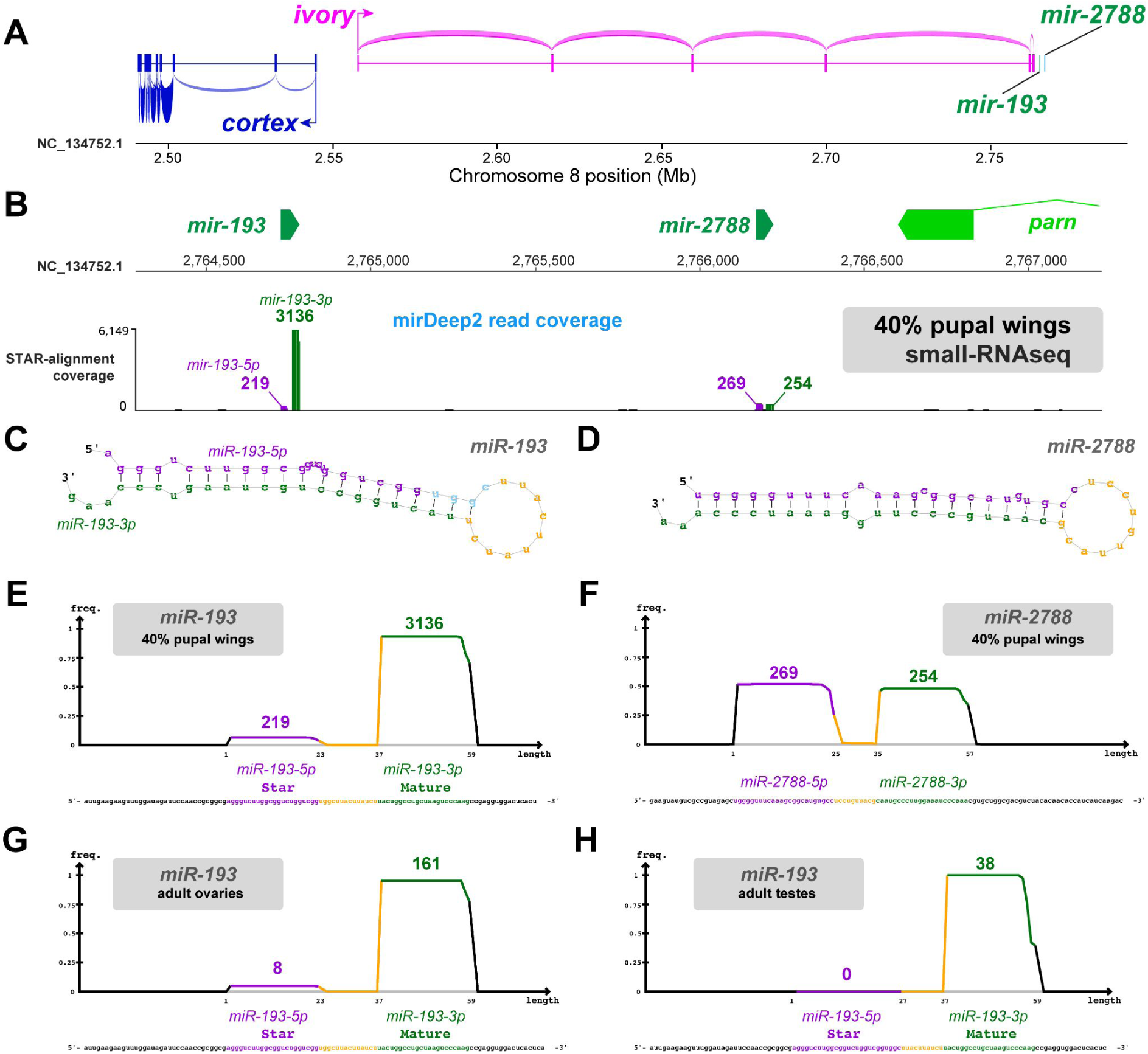
RNA-seq profiling of *cortex* and *ivory* and small RNA-seq profiling of mature *miR-193* and *miR-2788* RNAs. Related to Fig. 1. Short RNA-seq libraries from *A. gemmatalis* pupal wings at 40-80% of development were sequenced and mapped to the Dark reference genome using STAR. Separate, small RNA-seq libraries for *A. gemmatalis* pupal wings at the 40% stage, adult ovaries, and adult testes were also sequenced, and small-RNA reads were mapped to the genome using STAR and mirDeep2. **(A)** Sashimi plots showing splice support and coverage for *cortex* (blue) and *ivory* (magenta) expression. The positions of *mir-193* and *mir-2788* are also shown (green). **(B)** Coverage profiles of the *miR-193* and *miR-2788* RNAs, with strongest support for the *miR-193-3p* strand. Numbers in blue feature mirDeep2 read coverage. **(C-D)** Hairpin structures of predicted pre-miRNAs for *miR-193* and *miR-2788*. (**E-H)** Size distributions and relative abundances of Star and Mature miRNA processed transcripts predicted by mirDeep2 for *miR-193* in each sampled tissue (E, G, H), and for *miR-2788* in 40% pupal wings (F).

**Figure S6.**
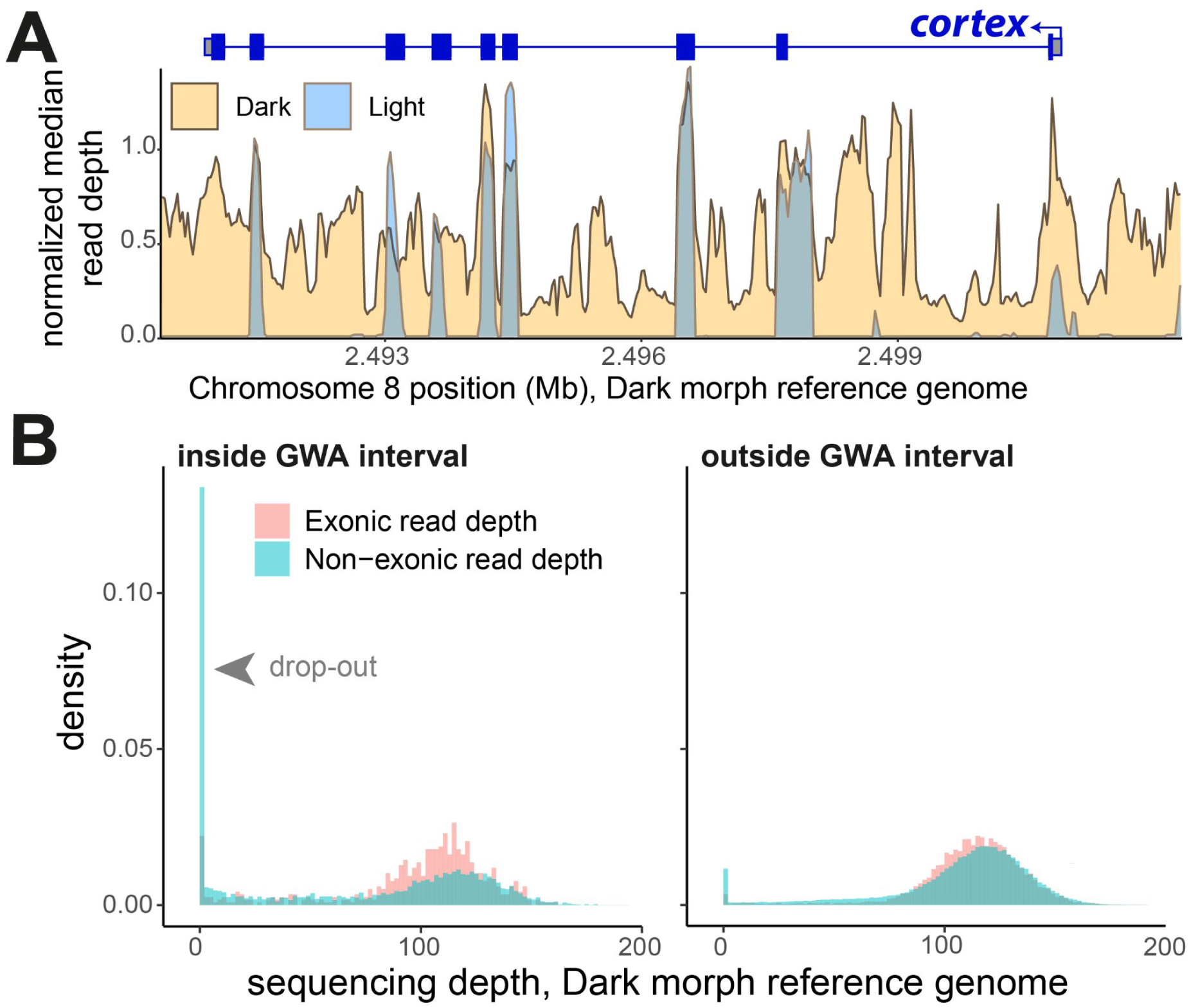
Coverage profiles for Dark and Light morphs across the *cortex* region and the GWA interval. Related to Fig. 2. **(A)** Normalized read depth profiles across the *cortex* gene, located 5’ of *ivory:mir-193*, using Pool-seq reads mapped to the Dark morph reference genome. Light-morph reads show reduced coverage across much of this region relative to Dark-morph reads, consistent with strong sequence divergence between morph-associated haplotypes. **(B)** Distributions of read depth in 100-bp windows for Light individuals, shown separately for exonic (red) and non-exonic (blue) regions inside and outside the GWA interval. Non-exonic regions within the GWA interval show pronounced read-mapping dropout relative to the genomic background.

**Figure S7.**
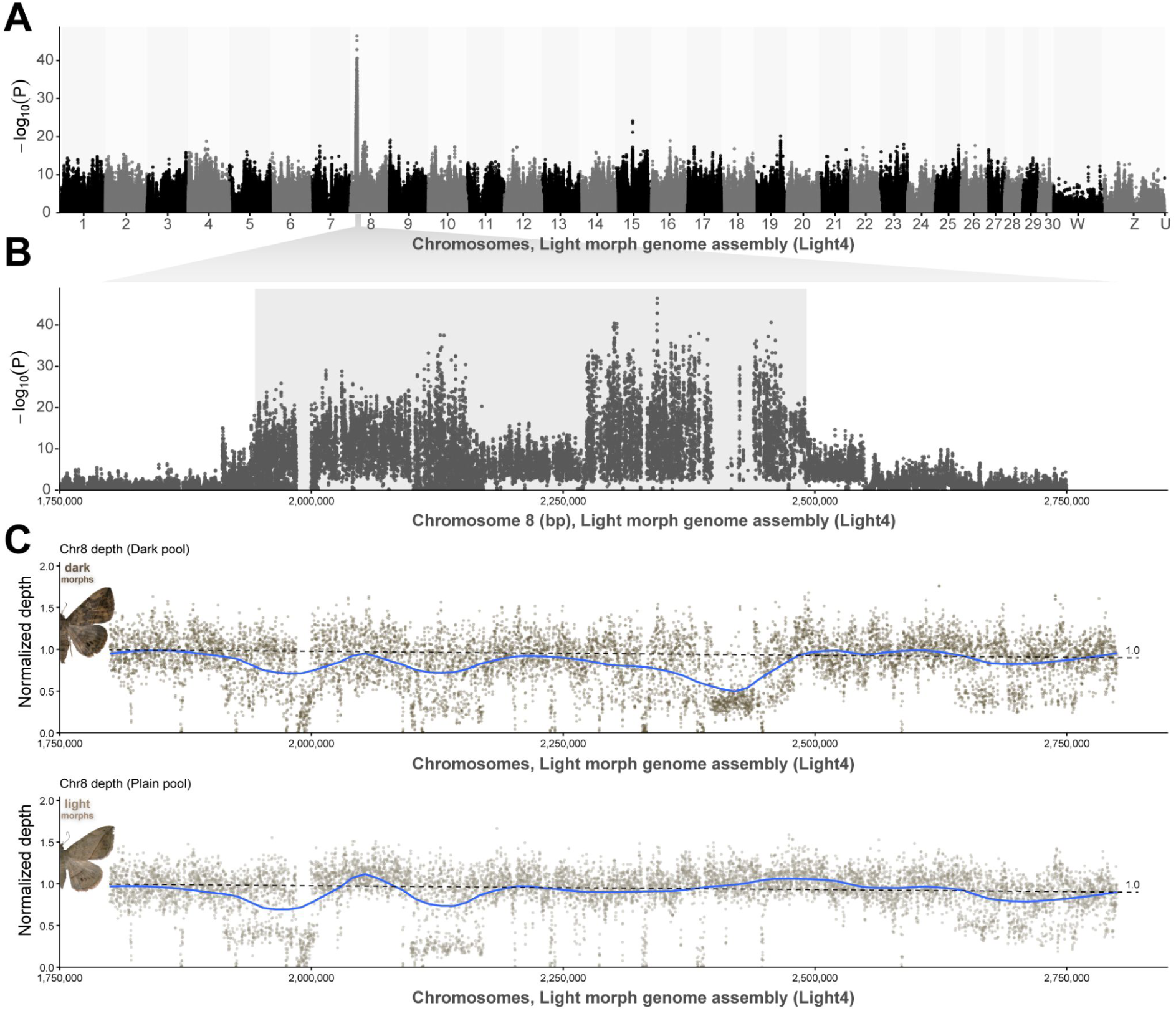
Pool-GWAS and coverage using a Light morph genome assembly as reference. Related to Fig. 1 and Fig. 2. **(A)** Genome-wide association scan for melanism based on pooled sequencing of Dark and Light morphs, with SNPs called against a Light morph assembly (Hifi pseudo-assembly of homozygous individual Light4). For each SNP, allele counts in the two pools were compared using Fisher’s exact test in PoPoolation2, and –log10(*P*) values are plotted by genomic position. **(B)** Expanded view of the associated region on chromosome 8. The association profile is similar to that shown in Figure 1. **(C)** Read-depth profiles for Dark and Light pooled libraries across the candidate interval using the Light morph reference genome.

**Figure S8.**
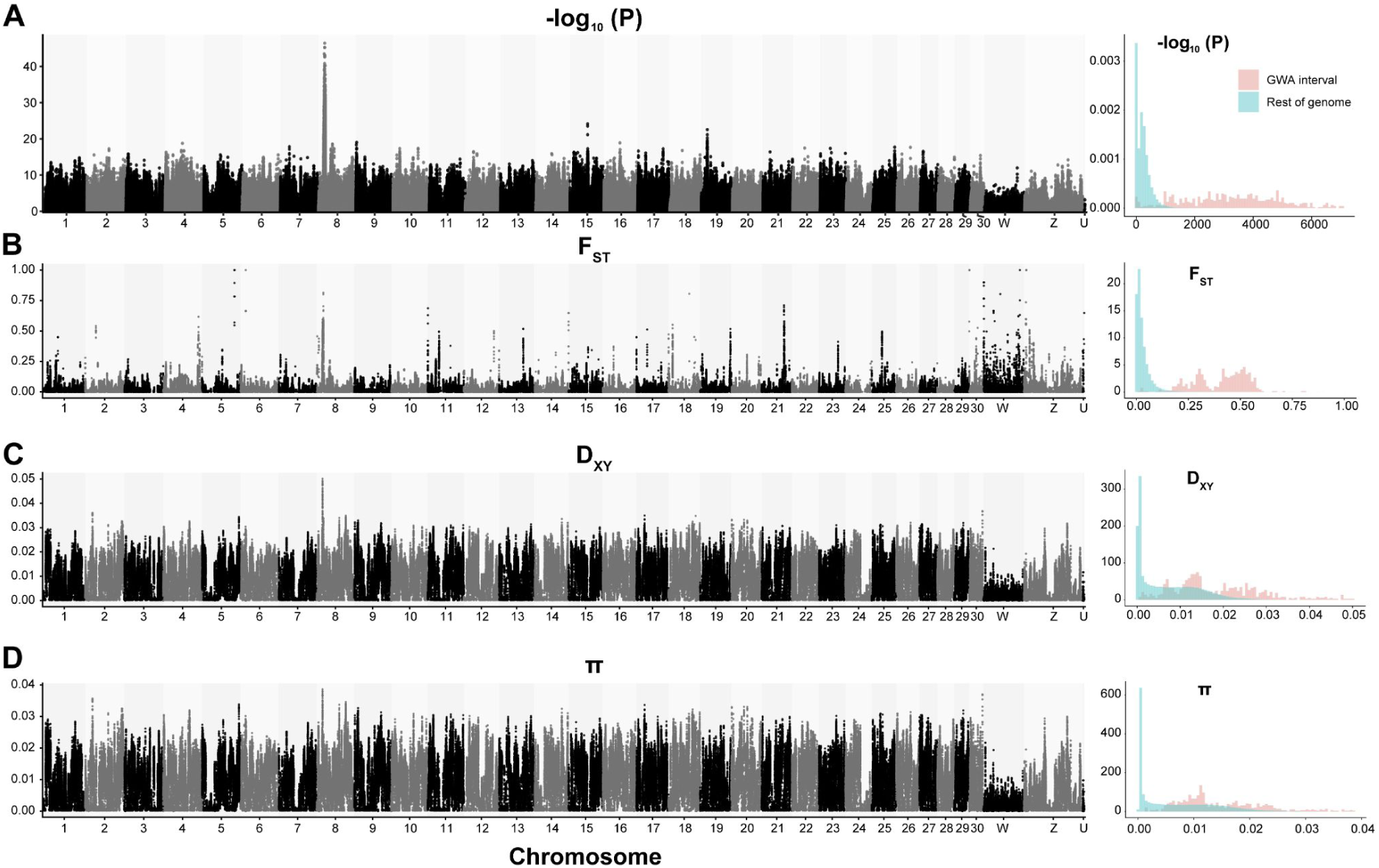
Genetic differentiation and nucleotide diversity across the whole genome and GWA interval. Related to Fig.1 and Fig.2. Each row shows a genome-wide scan of a summary statistic across the Dark reference genome (left) together with density distributions comparing values inside the GWA interval and across the rest of the genome (right). The combined pattern of high F_ST_, D_XY_, and π at the GWA melanic interval is consistent with the maintenance of deeply divergent haplotypes. **(A)** Fisher’s exact tests of association between pooled allele counts and phenotype (Dark versus Light), calculated in PoPoolation2. Because Pool-seq yields high read counts, Fisher’s exact test can produce extremely small *P* values; here, these values are shown as a ranking statistic rather than a precise estimate of effect size. **(B-C)** F_ST_ fixation index and D_XY_ absolute genetic divergence between the Dark and Light Pool-Seq groups. **(D)** Nucleotide diversity (π) calculated within all sequenced individuals.

**Figure S9.**
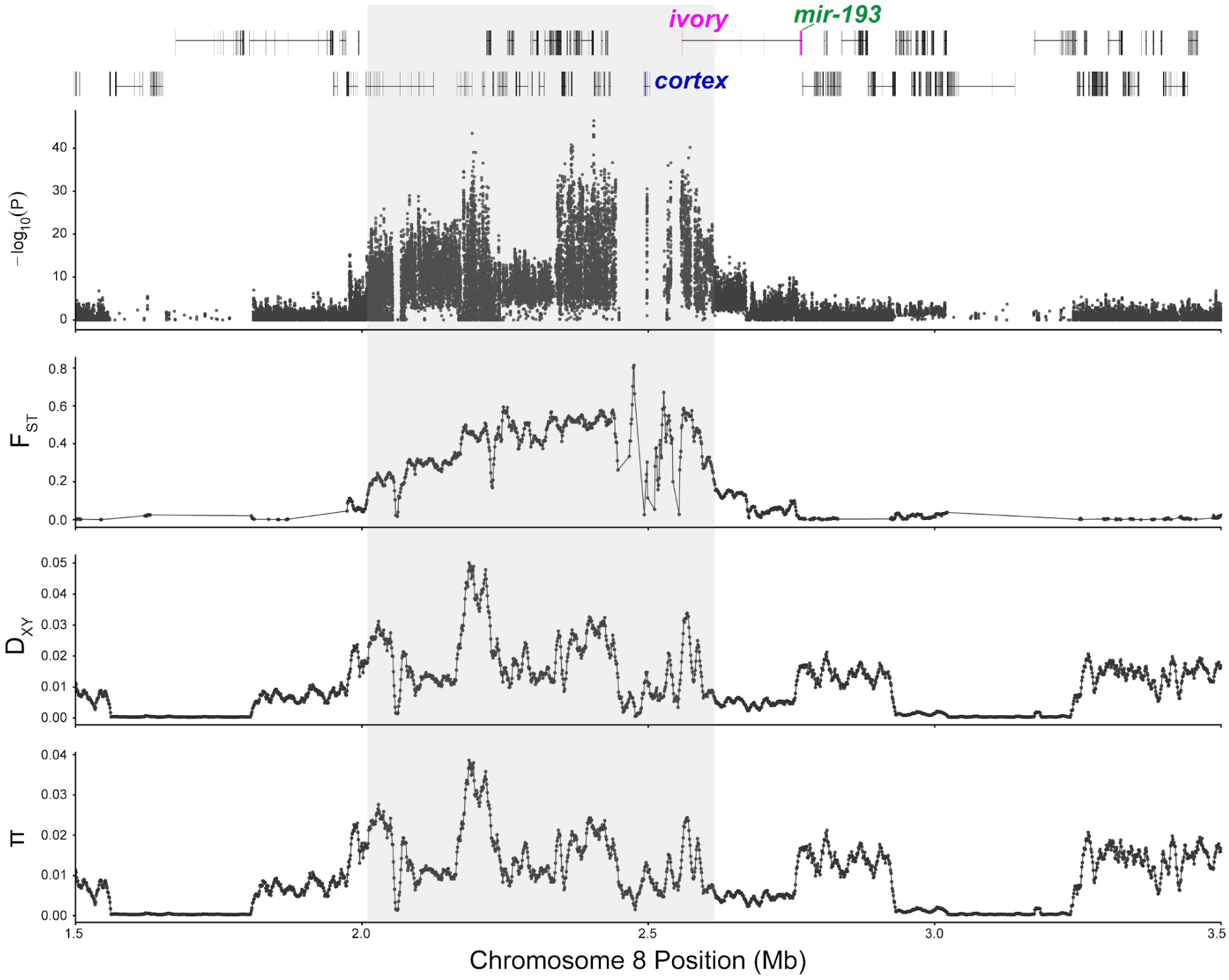
Elevated genetic differentiation and nucleotide diversity across the GWA interval. Related to Fig.2. Fisher’s exact test phenotype-association scores, F_ST_, D_XY_, and π measured on the Pool-Seq data as in Figure S8, are shown across the GWA interval in the Dark morph reference genome. Because some portions of the interval show read-mapping dropout, these statistics are likely underestimated in the most divergent regions.

**Figure S10.**
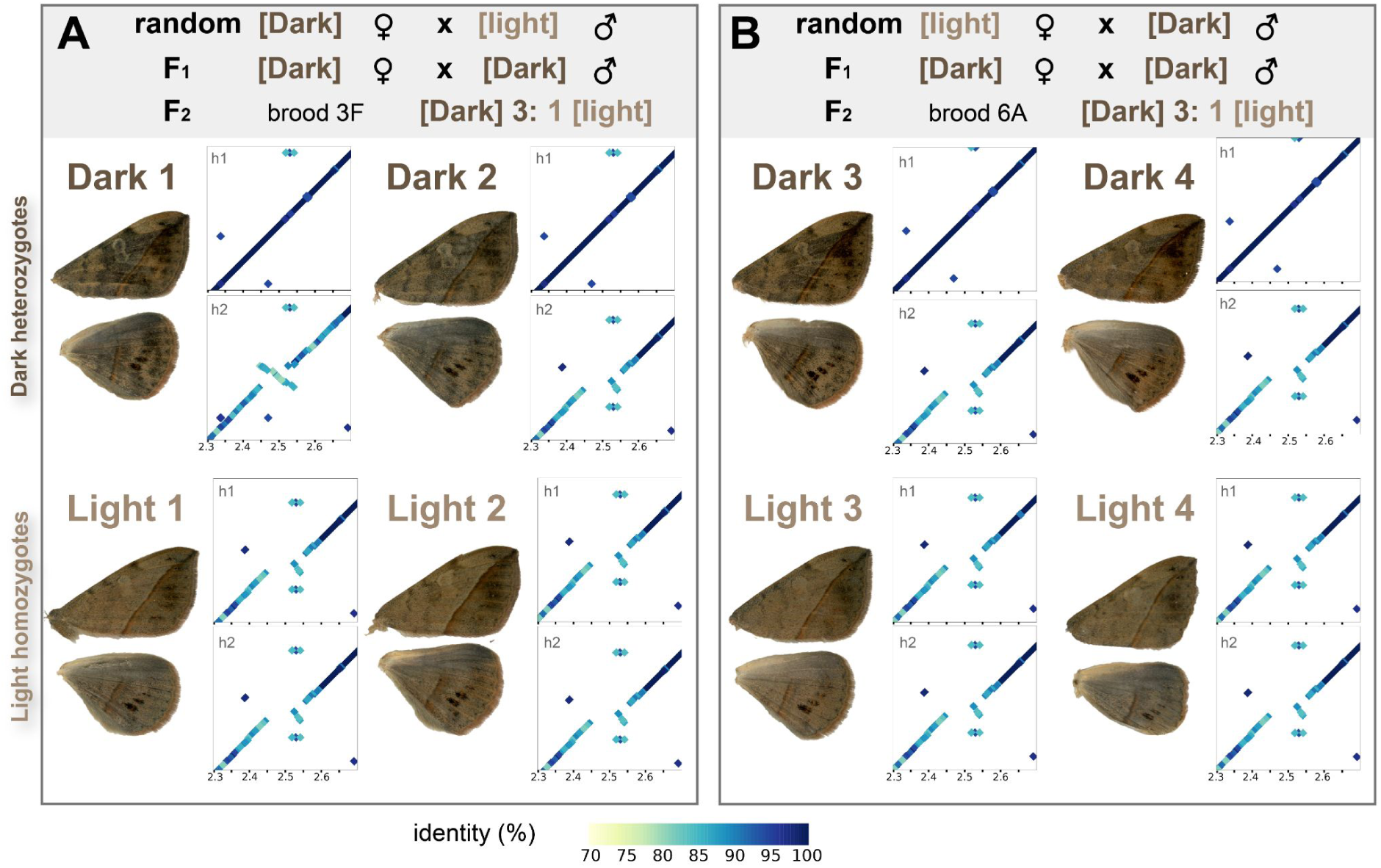
Dotplots of the SV region between the Dark reference haplotype against phased haplotypes. Related to Fig.2 and Fig.3. Representative forewing and hindwing images are shown for each PacBio HiFi–sequenced individual, together with dotplots comparing each phased haplotype (h1, h2) to the Dark reference assembly across the most divergent region of the morph-associated locus. Dot color indicates sequence identity percentage. Across the eight individuals, three haplotypes segregate in this interval. Haplotype A is largely collinear with the reference and found in all Dark individuals (shown as Dark h1 haplotypes here). Haplotype B was found in a single individual (Dark 1, h2) and shows a large inversion pattern. Haplotype C shows a translocated inversion pattern. It is found at a heterozygous state in 3 Dark individuals, and at the homozygous state in all Light individuals. Haplotypes A (Dark-dominant) and C (Light-recessive) are highly divergent, consistent with the presence of ancient structural haplotypes segregating at this locus.

**Figure S11.**
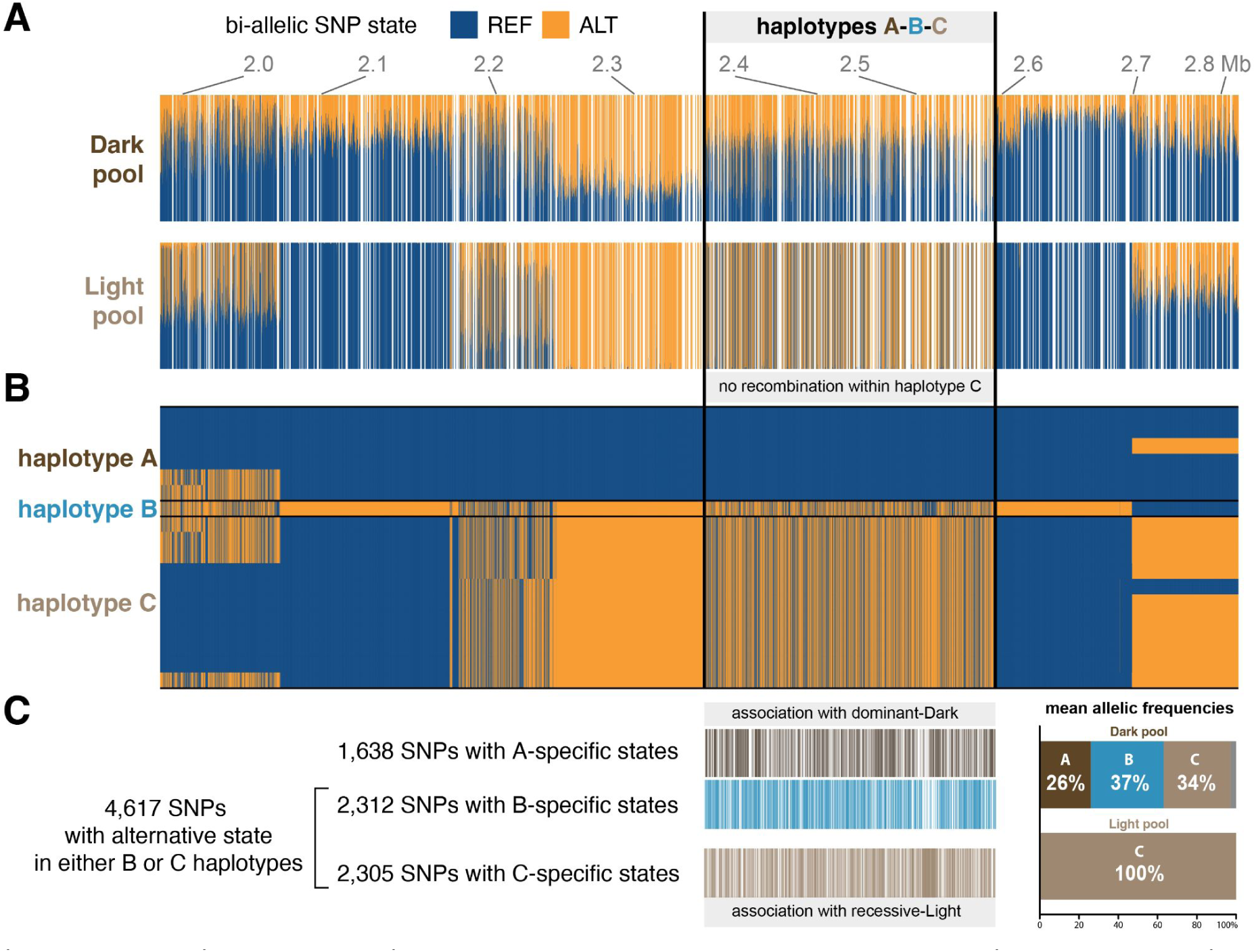
Allelic frequency differences follow a three haplotype structure in Dark versus Light Pool-Seq. Related to Fig 2. **(A)** Relative allele frequencies of biallelic SNPs within 25 Dark and 23 Light morphs sequenced in the Pool-Seq experiment. Blue features genotypes found in the Dark-homozygous reference genome. Orange corresponds to alternative genotypes at each biallelic SNP. Blanks correspond to variable sites that showed more than two alleles in the Pool-Seq experiment, but were sampled as biallelic states in the 8 phased genome assemblies. The C haplotype is at the homozygous state in the Light pool and showed no evidence of recombination in this outbred population. **(B)** Genotype plots of phased haploid assemblies for the Dark reference genome and eight additional individuals from two independent F_2_ families, sorted by haplotype in the A-B-C haplotype block. **(C)** Positions of the informative SNP sites for each haplotype. Histograms summarize the haplotype frequencies for each pool, as estimated from the mean allelic frequency for their informative sites (Fig. 2F).

**Figure S12.**
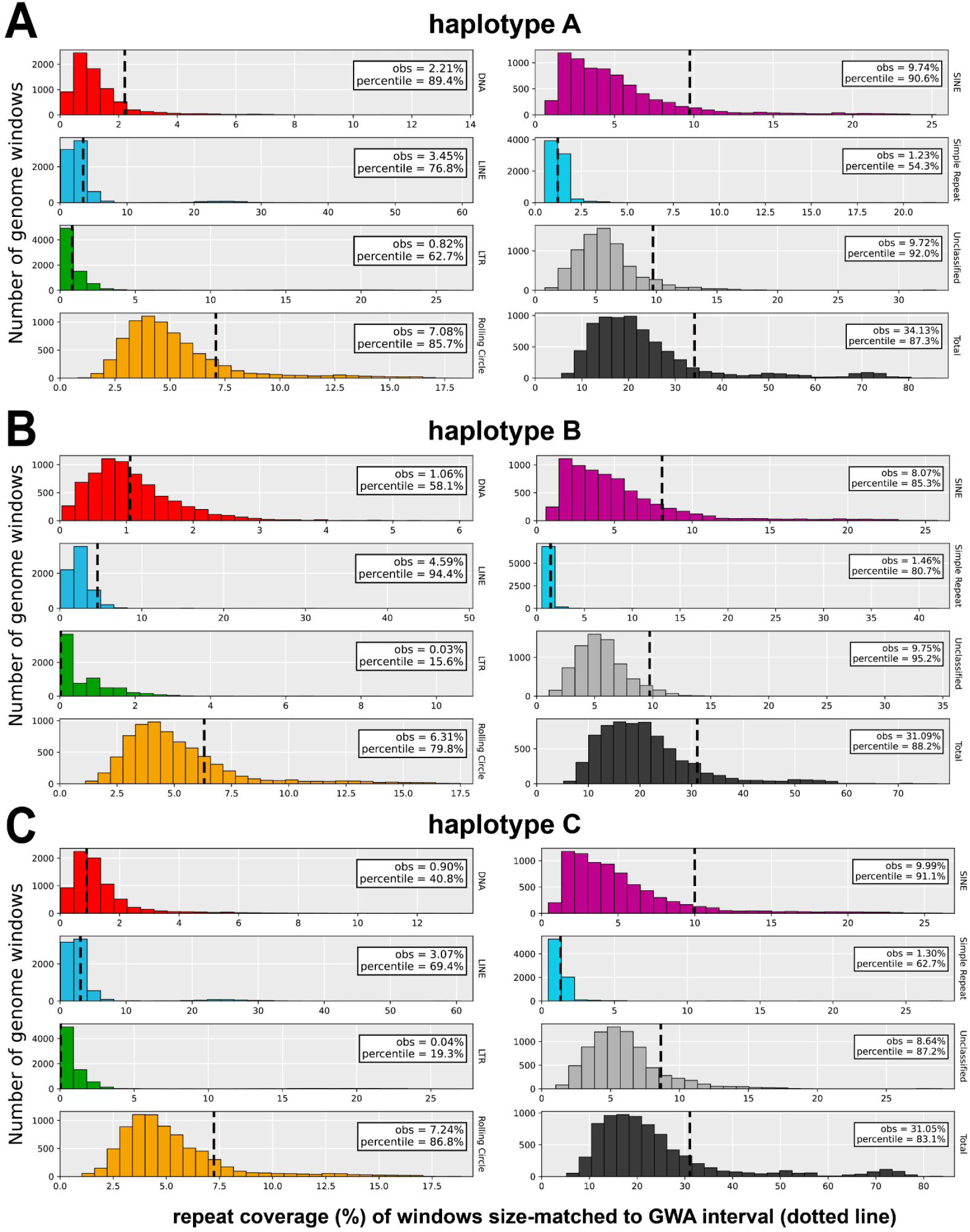
Repeat content of the chromosome 8 GWA interval relative to genome-wide windows of matched size. Related to Fig3. For each haplotype group (A–C), histograms show the genome-wide distribution of repeat-covered fraction in interval-sized windows for major Transposable Element and repeat categories and total repeat content. Dashed vertical lines indicate the observed repeat coverage of the hyperdivergent interval, and inset boxes report the observed value and its genome-wide percentile.

**Figure S13.**
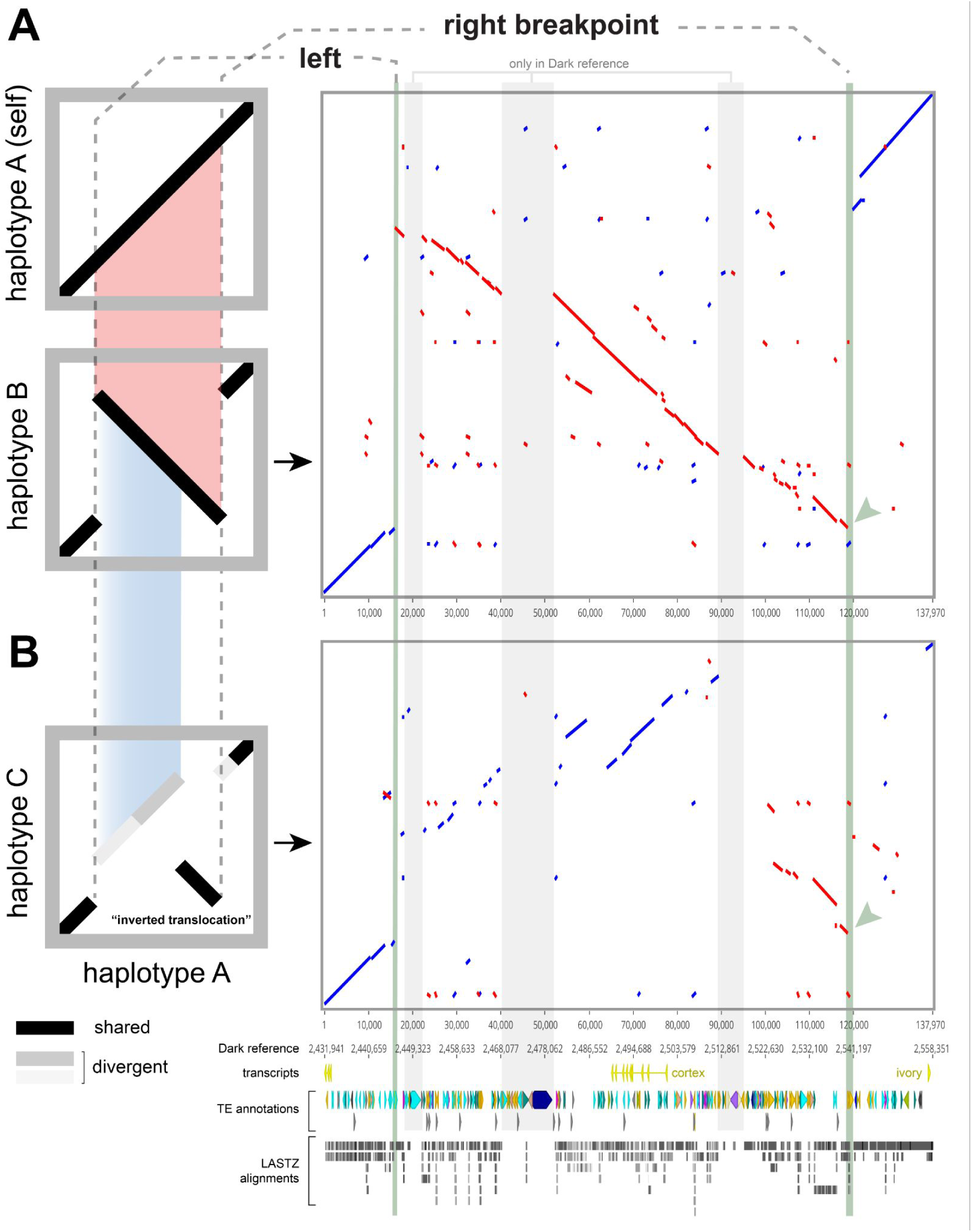
A double-inversion model for the inverted translocation observed between haplotypes A and C. Related to Fig.3. **Left:** schematic models on the left illustrate the theoretical dotplot relationships of two consecutive inversions that share a common right breakpoint. Regions of ectopic divergence, expected to reduce alignment scores, are shown in grey. **Right**: LASTZ dotplot alignments of haplotypes B and C relative to the haplotype A reference. Gene models, transposable-element annotations, and LASTZ alignments across the interval are shown below. **(A)** Haplotypes A and B differ by a large inversion. **(B)** Haplotypes A and C differ by an apparent inverted translocation. The right breakpoint is shared between the main SV variants (arrowheads) and nearly identical at the sequence level, supporting the double-inversion model.

**Figure S14.**
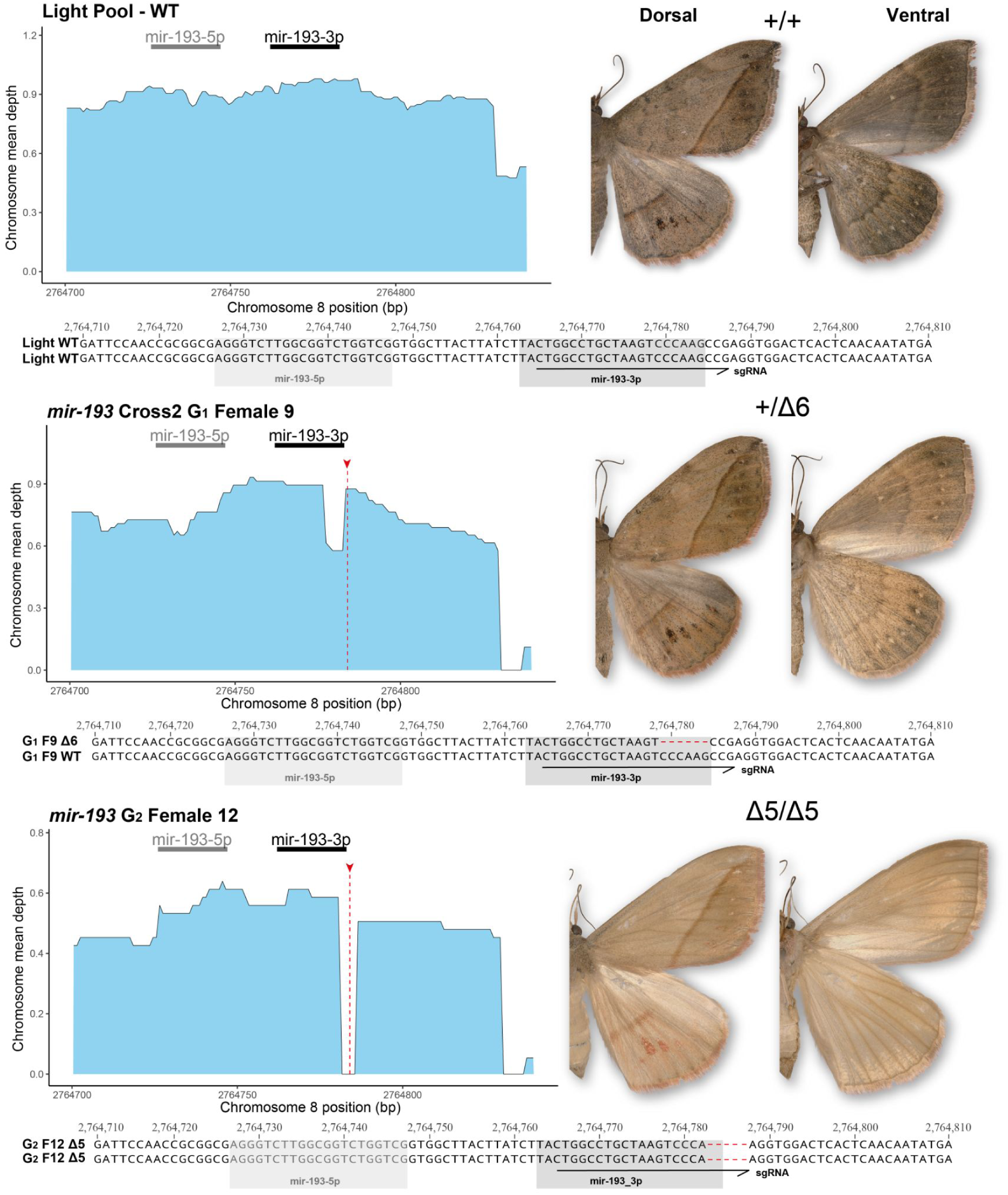
Whole genome sequence genotyping of *mir-193* CRISPR KOs. Related to Fig. 5. Mean whole-genome sequencing depth across the *mir-193* locus for wild-type Light pool (top), a G_1_ heterozygous *mir-193* crispant (middle), and a G_2_ homozygous crispant (bottom). Coverage from short-read data is plotted across the region spanning the annotated *mir-193 5p* and *3p* arms (black bars). In CRISPR genotyped individuals, sequencing depth drops locally at the expected sgRNA cut site (red dashed line), consistent with a deletion disrupting read mapping at the target. For each individual, photographs of the same moth are shown to the right (dorsal and ventral wing surfaces). Below each depth trace, both haplotypes are shown as aligned allele sequences (WT and/or mutant); mutant alleles contain the CRISPR-induced deletions (red).

**Figure S15.**
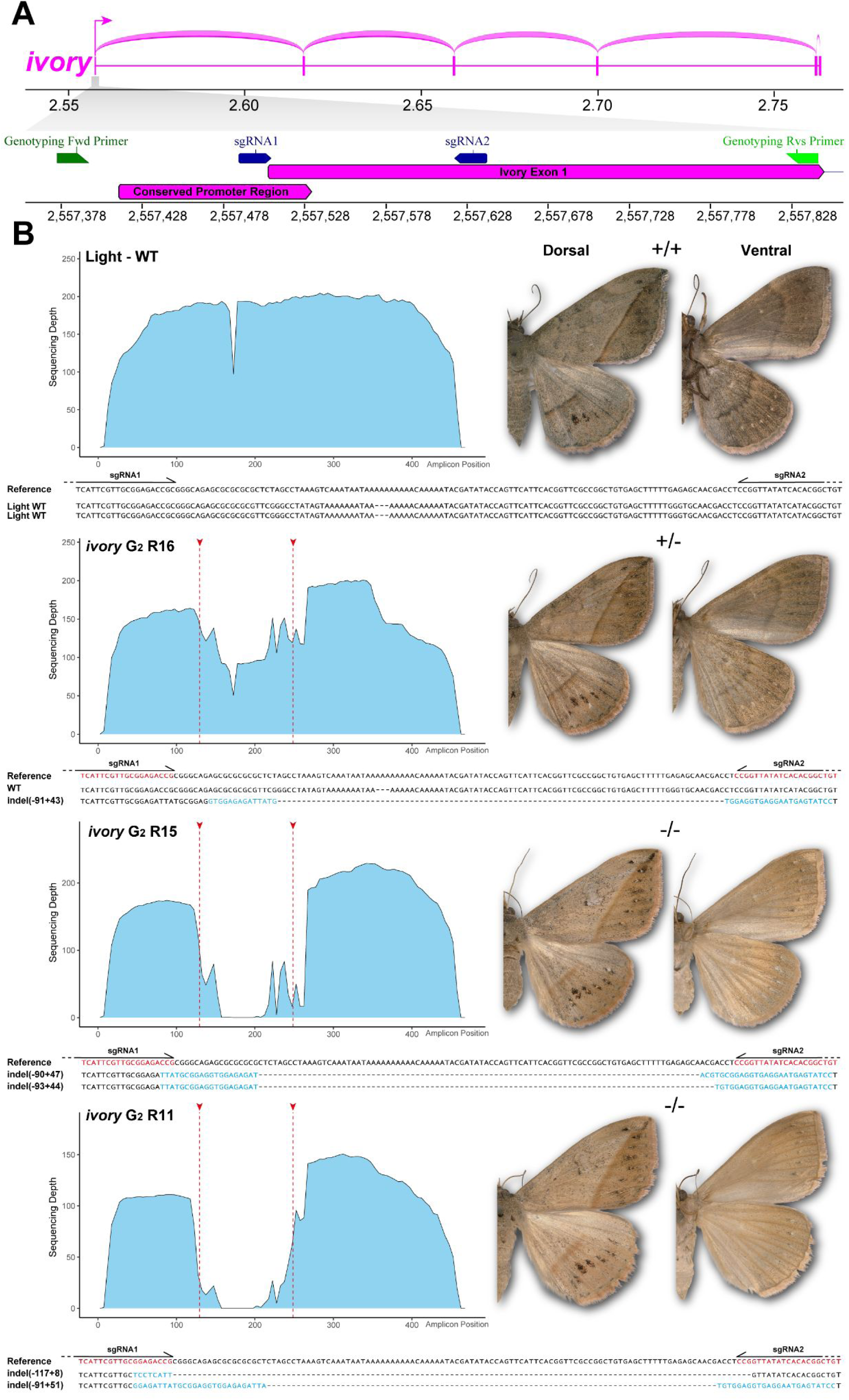
Long-read amplicon genotyping of *ivory* CRISPR KOs. Related to Fig. 5. **(A)** Sashimi plot showing the *ivory* exon structure above a zoomed-in annotation of the promoter/exon 1 region, with the positions of the sgRNAs and genotyping primers highlighted. **(B)** Long-read sequencing depth across *ivory* amplicons is shown for a wild-type Light individual (+/+), a G_2_ heterozygous *ivory* crispant (+/−), and two G_2_ compound heterozygous crispants (−/−). In CRISPR-genotyped individuals, sequencing depth drops locally at the expected sgRNA cut sites (red dashed lines), consistent with indels at the sgRNA target sites. For each genotyped individual, photographs of the corresponding moth are shown to the right (dorsal and ventral wing surfaces). Both haplotypes are shown as aligned allele sequences to the reference genome (WT and/or mutant). Mutations are highlighted as deletions (gaps/dashes in the alignment) and non-templated inserted sequence (blue), and each indel allele is labeled by its net deleted/inserted length.

**Table S1.**
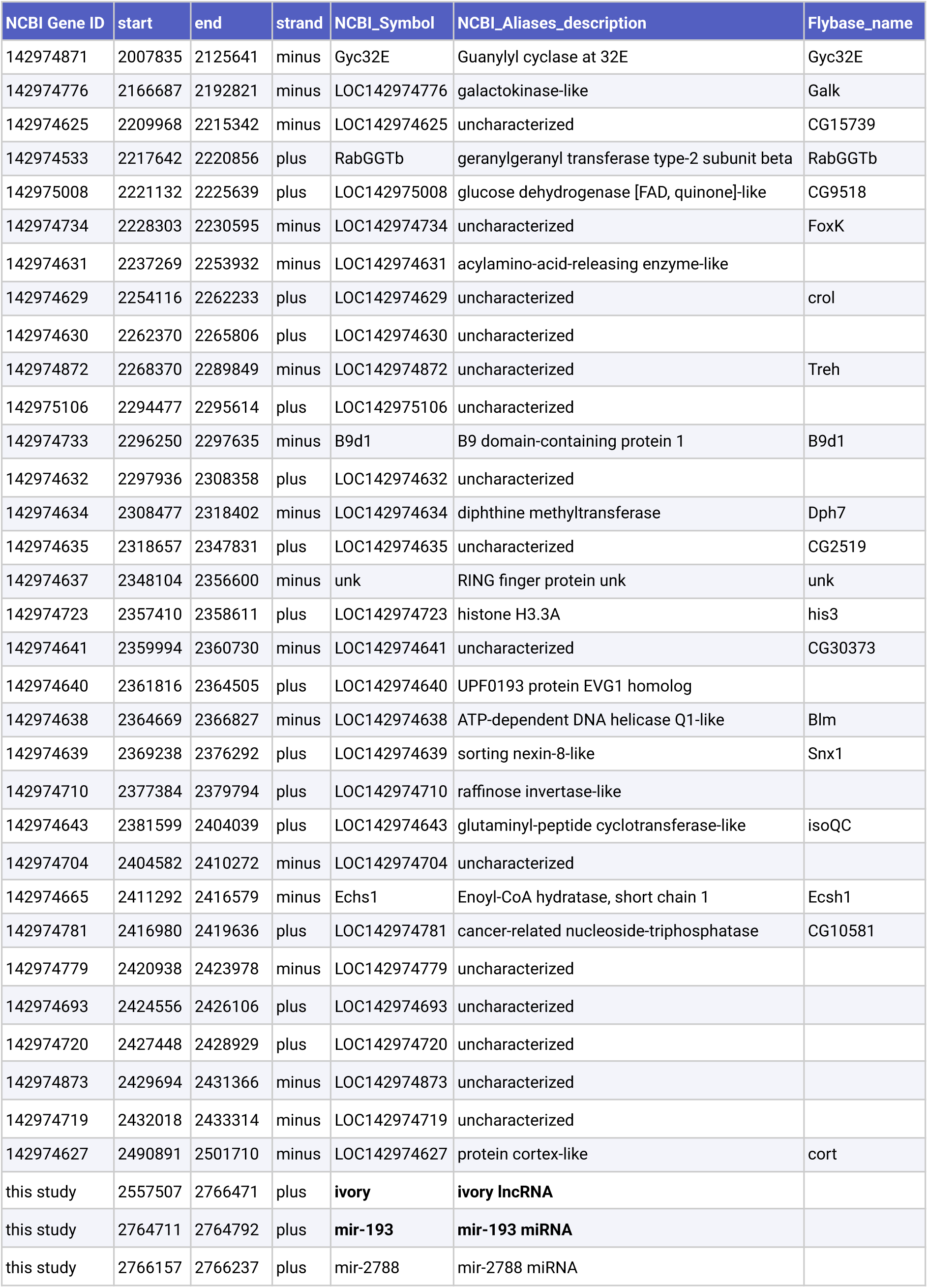
Gene annotations in the Pool-GWA interval. Positions are relative to the NCBI RefSeq genome chromosome 8 (NC_134752.1). Annotations of the *ivory* lncRNA, *mir-193* miRNA and *mir-2788* miRNA do not feature on the online RefSeq annotation, and among them, only the first exon of the ivory lncRNA is included in the GWA interval.

**Table S2.** Summary of CRISPR injection experiments. Related to Fig. 4 and STAR Methods.

| sgRNA(s) | Inj. Embryos (Ninj) | L3 Survival (Nhat) | Pupae | Adults (Nadu) | Crispant (Nmut) | Inj. time (min AEL) | Cas9:sgRNA (ng/μl) | Hatching Rate (Nhat/Ninj) | Penetrance (Nmut/Nadu) |
| --- | --- | --- | --- | --- | --- | --- | --- | --- | --- |
| *Ag_mir-193* | 973 | 194 | 170 | 153 | 126 | 45-129 | 500:125:125 | 19.94% | 82.35% |
| *Ag_ivory1+2* | 989 | 27 | 20 | 19 | 17 | 27-120 | 500:125:125 | 2.73% | 89.47% |

**Table S3.**
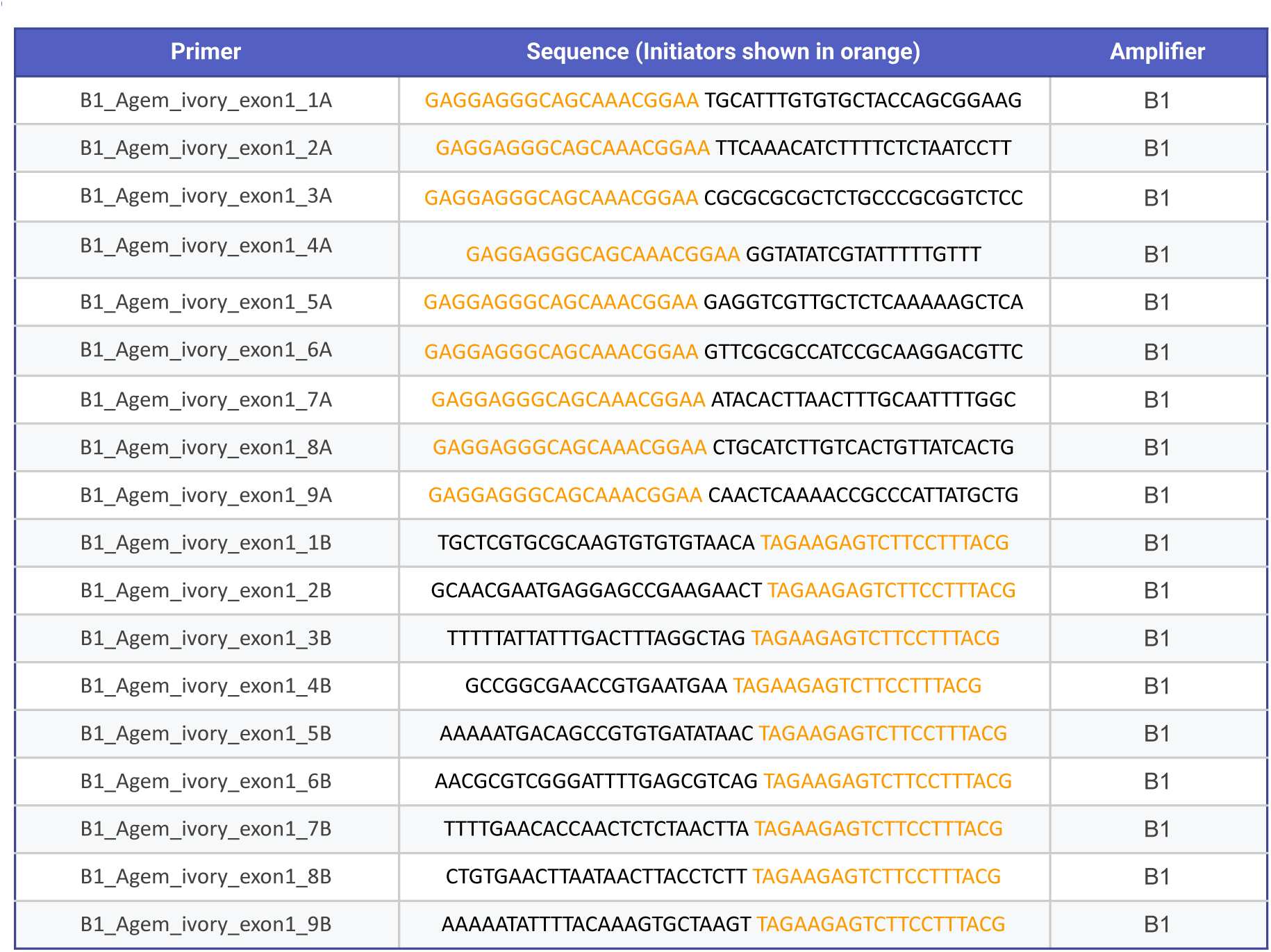
List of oligonucleotides used for the generation of *ivory* HCR probes. Related to STAR Methods and Figure 3.

**Table S4.**
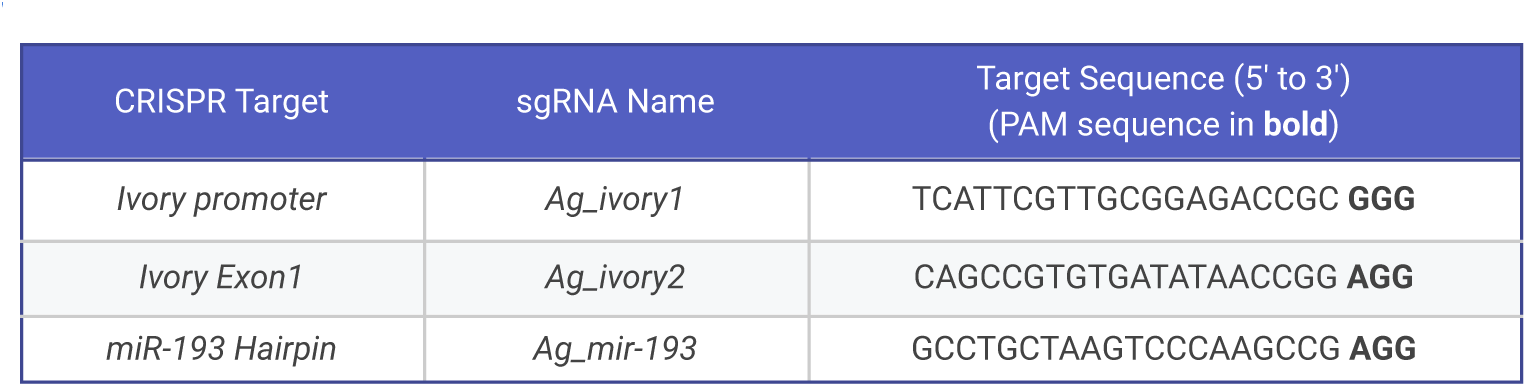
sgRNA targets used in CRISPR mKO experiments. Related to Fig. 4.

