## Supplementary Files for "Hyperdivergent haplotypes control melanic camouflage in a polymorphic moth"

| REAGENT or RESOURCE | SOURCE | IDENTIFIER |
|---|---|---|
| **Chemicals, peptides, and recombinant proteins** | | |
| Cas9-2xNLS protein | QB3 Macrolab, UC Berkeley | N/A |
| Bovine Serum Albumin (BSA) | ThermoFisher Scientific | Cat#J65788.09 |
| 4',6-diamidino-2-phenylindole, dihydrochloride (DAPI) | ThermoFisher Scientific | Cat#D1306 |
| EDTA, 0.5 M, pH 8.0 | Bioland Scientific | Cat#EDTA01 |
| EGTA, 0.5 M, pH 8.0 | bioWorld | Cat#40520008-1 |
| Formaldehyde, 37% | Sigma Aldrich | Cat#252549-500ML |
| Glycerol, ≥99.0% | Sigma Aldrich | Cat#G5516 |
| Low EDTA TE (1x) buffer, pH 8.0 | Quality Biological | Cat#351-324-721 |
| Phosphate Buffer Saline (PBS), 10X | UFC Bio | Cat#BPBS74-10X |
| Saline-sodium citrate (SSC), 20X | Quality Biological | Cat#351-003-101 |
| Triton X-100 | Sigma Aldrich | Cat#9036-19-5 |
| Tween20 | Bioworld | Cat#42030016-1 |
| Benzalkonium Chloride solution, 50% | Sigma-Aldrich | Cat#63249-500ML |
| Phenol Red, 0.5 % | Sigma-Aldrich | Cat#P0290-100ML |
| TRI Reagent | Zymo Research | Cat#R2050 |
| Proteinase K | Promega | MC5005 |
| SSC, 20x | Any | N/A |
| Methanol | Any | N/A |
| Glycine, 10 g/L | Any | N/A |
| **Critical Commercial Assays** | | |
| Taq 2x Master Mix | New England Biolabs | M0270 |
| Luna Universal qPCR Master Mix | New England Biolabs | Cat#M3003 |
| PrimeScript RT Reagent Kit with gDNA Eraser | TaKaRa Bio | Cat#RR047A |
| AccuGreen Broad Range dsDNA Quantitation Kit | Biotium Inc. | Cat#31069 |
| DNA Clean & Concentrator-5 | Zymo Research | Cat#D4004 |
| Quick-DNA Miniprep Plus Kit | Zymo Research | Cat#D4068 |
| Direct-zol RNA Miniprep Kit | Zymo Research | Cat#R2051 |
| HCR v.3.0 Alexa Fluor 647 Amplifier (B1) | Molecular Instruments | N/A |
| HCR v3.0 Hybridization Buffer, Probe Wash Buffer, Amplification Buffer | Molecular Instruments | N/A |
| **Deposited Data** | | |
| *ilAntGemm2* RefSeq genome assembly and annotation | This study | NCBI: GCF_050436995.1 |
| *ilAntGemm2* genome assembly, alternate haplotype | This study | NCBI: GCA_050436975.1 |
| *A. gemmatalis,* RNAseq raw reads | This study | NCBI: PRJNA1165401 |
| *A. gemmatalis,* PoolSeq raw reads for Dark vs. Plain phenotypes | This study | NCBI: PRJNA1338508 |
| **Experimental models: Organisms/strains** | | |
| *Anticarsia gemmatalis,* Stoneville strain | Benzon Research | NCBI:txid129554 |
| **Oligonucleotides** | | |
| Oligonucleotides for HCR, see Tables S1 to S3 | This study | N/A |
| **Software and algorithms** | | |
| C++ / Adobe Illustrator and Adobe Photoshop | Adobe | URL: https://www.adobe.com/ |
| C/C++ / Blast+ v.2.16.0+ | Camacho et al. 2009 | URL: https://blast.ncbi.nlm.nih.gov/doc/blast-help/downloadblastdata.html |
| Java / FastQC v.0.12.1 | GitHub/ S-andrews | URL: https://github.com/s-andrews/FastQC |
| Java / FIJI | Schindelin et al. 2012 | URL: https://imagej.net/software/fiji/downloads |
| C++ / Flexbar v.3.5.0 | Roehr et al. 2017 | URL: https://github.com/seqan/flexbar |
| Java / Geneious Prime | Geneious | URL: https://www.geneious.com/ |
| Java / Integrative Genomics Viewer (IGV) v.2.19.2 | Thorvaldsdóttir et al. 2013 | URL: https://igv.org/doc/desktop/ |
| Python / Lep_BUSCO_Painter v.1.0.0 | Wright et al. 2024 | URL: https://github.com/charlottewright/lep_busco_painter |
| C / MAFFT | Katoh & Standley 2013 | URL: https://www.geneious.com/plugins/mafft |
| Perl / miRDeep2 | Friedländer et al. 2012 | URL: https://github.com/rajewsky-lab/mirdeep2 |
| Python / Napari | Chiu et al. 2022 | URL: https://github.com/napari/napari |
| C / Samtools v.1.2.2 | Danecek et al. 2021 | URL: https://github.com/samtools/samtools |
| C/C++ / STAR v.2.7.11b | Dobin et al. 2016 | URL: https://github.com/alexdobin/STAR |
| Nim / Mosdepth v0.3.3 | Pedersen and Quinlan, 2018 | URL: https://github.com/brentp/mosdepth |
| C/C++ Minimap2 v2.2.30 | Li, 2018 | URL: https://github.com/lh3/minimap2 |
| Python / Insitu Probe Generator v.0.3.2 | Github/Ryan Null | URL: https://github.com/rwnull/insitu_probe_generator |
| Perl / Popoolation2 v1.201 | Kofler et al., 2011 | URL: https://github.com/lczech/popoolation2 |
| C / BWA (Burrows-Wheeler Aligner) v.0.7.17-r1188 | Li and Durbin, 2010 | URL: https://github.com/lh3/bwa |
| C++ / Hifiasm v0.25.0-r726 | Cheng et al. 2021 | URL: https://github.com/chhylp123/hifiasm |
| Python / RagTag | Alonge et al. 2022 | URL: https://github.com/malonge/RagTag |
| Python / Liftoff | Shumate and Salzberg 2021 | URL: https://github.com/agshumate/Liftoff |

| | | | |
|---|---|---|---|
| | C / BCFtools | Danecek et al. 2021 | URL: https://samtools.github.io/bcftools/ |
| | C/C++ / BEDTools | Quinlan and Hall 2010 | URL: https://github.com/arq5x/bedtools2 |
| | C/C++ / MUMmer v4.0.1 | Marcais et al. 2018 | URL: https://mummer4.github.io/ |
| | C / LASTZ | Harris 2007 | URL: https://lastz.github.io/lastz/ |
| | C++ / grenedalf v0.6.3 | Czech et al. 2024 | URL: https://github.com/lczech/grenedalf |
| | Python / blastn2dotplots | Okuno et al. 2025 | URL: https://github.com/mokuno3430/blastn2dotplots |
| | R / GenotypePlot v0.2.1 | GitHub/Jim Whiting | URL: https://github.com/JimWhiting91/genotype_plot |
| | Pipeline / EarlGrey v7.2.1 | Baril et al. 2024 | URL: https://github.com/TobyBaril/EarlGrey |
| | Perl / RepeatMasker v4.1.7/v4.2.2 | Smit, Hubley and Green; Tempel 2012 | URL: https://www.repeatmasker.org/ |
| | Perl / RepeatModeler2 | Flynn et al. 2020 | URL: https://github.com/Dfam-consortium/RepeatModeler |
| | Database / Dfam and FamDB | Storer et al. 2021 | URL: https://www.dfam.org/ |
| | R / pegas package | Paradis 2010 | URL: https://cran.r-project.org/package=pegas |
| | R / ape package | Paradis and Schliep 2019 | URL: https://cran.r-project.org/package=ape |
| | R / vcfR package | Knaus and Grunwald 2017 | URL: https://cran.r-project.org/package=vcfR |
| | R / ggplot2 package v.3.4.2 | Tidyverse/Hadley Wickham | URL: https://ggplot2.tidyverse.org |
| | R / tidyverse package v.2.0.0 | Tidyverrse/Hadley Wickham et al. | URL: https://www.tidyverse.org/ |
| | R / cowplot package v1.1.3 | Wilke, 2025 | URL: https://CRAN.R-project.org/package=cowplot |
| | R / dplyr package v1.1.4 | Tidyverrse/Hadley Wickham et al. | URL: https://www.tidyverse.org/ |
| | R / readr package v2.1.5 | Tidyverrse/Hadley Wickham et al. | URL: https://www.tidyverse.org/ |
| | R / ggrastr package v1.0.2 | Viktor Petukhov et al. 2021 | URL: https://github.com/VPetukhov/ggrastr |
| | R / recolorize package v0.2.0 | Weller, 20205 | URL: https://hiweller.github.io/recolorize/ |
| | **Other** | | |
| | Black Enameled Pins, Size 1 | Pin-It Entomological Supply | N/A |
| | Copper Wire Mesh,100 x 100 Mesh, 0.0045" Diameter Wire | Small Parts | Cat#CU-100-0045-01 |
| | LocknLock Rectangular, 1L | LocknLock | Cat#67140 |
| | Small Dissection Petri Dish, Clear, 50 mm Dia x 17 mm H | Living Systems Inc | Cat#DD-50-S-3PK |
| | Stainless Steel Cup Holder | Da Vinci | Cat#B06W2JBLJJ |
| | Steel Woven Wire Cloth Disc, 40 X 40 Mesh, 2-9/16" Diameter | McMaster-Carr | Cat#2812T43 |
| | Vannas Spring Scissors - 2.5 mm | Fine Science Tools | Cat#15000-08 |
| | Borosilicate capillaries with filament | WorldPrecision Instruments | Cat#18100F-3 |
| | Gravity needle puller, PC-10 | Narishige International | N/A |
| | Three-axis MM33 right-handed manipulator | Drummond Scientific | Cat#3-000-024-R |
| | Single pressure micro-injectors with footswitch | Tritech Research Inc. | #CatMINJ-1 |
| | Pulse-length control module | Tritech Research Inc. | Cat#MINJ-2 |
| | Needle holder | Tritech Research Inc. | Cat#MINJ-4 |
| | Polyurethane tubing 1/4” OD, 1/8” ID | Tritech Research Inc. | Cat#TT-1-4OD |
| | 3-way Tee air splitter | Tritech Research Inc. | Cat#MINJ-3TQC |
| | Binocular stereomicroscopes with 10–25× magnification | Any | N/A |
| | Cardboard Freezer Box, with 196 place PCR inserts | Argos Technologies | Cat#FBZ-1196W |
| | Disposable Sterile Scalpels with Plastic Handle, Pack of 10 | JMU | UPC#840456203194 |
| | Extra Strength Microgreen and Seedling Trays 10" x 10" | Bootstrap Farmer Store | UPC#811284031754 |
| | Multiple Species Diet | Southland Inc. | N/A |
| | Plastic Pestle and 1.5ml Tube | Bel-Art | F19923-0000 |
| | Qubit 3.0 fluorometer | Thermo Fischer Scientific | Cat#Q33216 |
